# NKX2-1 controls lung cancer progression by inducing DUSP6 to dampen ERK activity

**DOI:** 10.1101/2021.03.04.433941

**Authors:** Kelley Ingram, Shiela C. Samson, Rediet Zewdu, Rebecca G. Zitnay, Eric L. Snyder, Michelle C. Mendoza

## Abstract

Lung cancer remains a leading cause of cancer death, with unclear mechanisms driving the transition to aggressive cancer with poor prognosis. The RAS→RAF→MEK→ERK pathway is hyper-activated in ∼50% of human lung adenocarcinoma (LUAD). An initial activating mutation induces homeostatic feedback mechanisms that limit ERK activity. Additional, undefined genetic hits overcome the feedback, leading to high ERK activity that drives malignant progression. At detection, the majority of LUADs express the homeobox transcription factor NKX2-1, which also limits malignant progression. NKX2-1 constrains LUAD, in part, by maintaining a well-differentiated state with features of pulmonary identity. We asked if loss of NKX2-1 might also contribute to the release of ERK activity that drives tumor progression. Using human tissue samples and cell lines, xenografts, and genetic mouse models, we show that NKX2-1 induces the ERK phosphatase DUSP6. In tumor cells from late-stage LUAD with silenced *NKX2-1*, re-introduction of NKX2-1 induces DUSP6 and inhibits cell proliferation and migration and tumor growth and metastasis. CRISPR knockout studies show that DUSP6 is necessary for NKX2-1-mediated inhibition of tumor progression *in vivo*. Further, DUSP6 expression is sufficient to inhibit RAS-driven LUAD. We conclude that *NKX2-1* silencing, and thereby *DUSP6* downregulation, is a mechanism by which early LUAD can unleash ERK hyperactivation for tumor progression.

## Introduction

Lung cancer and specifically lung adenocarcinoma (LUAD) remains the leading cause of cancer death world-wide. New therapies that molecularly target driver mutations have improved patient outcome. However, the development of resistance mutations necessitate long-term, dynamic, and sequential cycles of targeted therapeutics to combat resistance (1). Better understanding of the molecular mechanisms of LUAD progression will enable the development of new therapeutic strategies for long-term patient management.

The mitogen-activated protein (MAP) kinase extracellular signal-regulated kinase (ERK) is an established initiator and driver of lung adenocarcinoma (LUAD). ERK is activated downstream of receptor tyrosine kinase signaling to the RAS→RAF→MEK→ERK pathway. Nearly 50% of LUADs harbor mutations in *RAS* genes, *BRAF*, or *NF1*, which encodes the RAS GTPase activating protein (2, 3). An additional 13% harbor mutations upstream receptor tyrosine kinases *EGFR*, *ERBB2*, *MET*, *ALK*, *RET*, and *ROS1* (2, 3). Genetically-engineered mouse models show that these mutations are sufficient to initiate pre-cancerous and low-grade lesions (4, 5). Tumor initiation and maintenance requires ERK activity, as MEK inhibition or ERK deletion blocks tumor development (4, 6, 7). Yet, the initial ERK activation is insufficient for progression to metastatic cancer (5, 8).

Active, phosphorylated ERK (phospho-ERK) is associated with more aggressive LUAD (9, 10), suggesting that ERK promotes progression. This premise is also supported by experiments in mouse models. When *Kras* mutation is combined with *Trp53* or *Lkb1* deletion and given time for the spontaneous acquisition of additional mutations, high-grade metastatic cancer develops (4, 8, 11–14). These aggressive cancers are associated with an increase in phospho-ERK (11, 12). Increased ERK signaling is sufficient for progression. Paradoxical CRAF activation, by inhibition of BRAF^V600E^ in the context of KRAS activation, stimulates ERK activity and accelerates tumor progression (15–17). How the initial low level of ERK activity transitions to high activity sufficient for driving progression to metastasis is unclear. Plausible mechanisms include inactivation of the wildtype *KRAS* allele and loss of the ERK pathway negative feedback loops (12, 18–20).

LUAD presents with heterogeneous histopathology and differentiation states (21). In addition to exhibiting high ERK activity, the most aggresive cancers are also poorly differentiated (22, 23). Mouse models of LUAD suggest that de-differentiation involves transition through a state of latent gastric differentiation, which is initiated by loss of the lineage transcription factor NKX2-1/TTF-1 (24–26). In KRAS^G12D^-driven tumors, *NKX2-1* deletion increases phospho-ERK and accelerates progression to invasive carcinoma and metastasis (8, 24, 27). Clinically, low *NKX2-1* expression portends a poor prognosis (28–30). We reasoned that in addition to beginning the de-differentiation process, a repressed-*NKX2-1* status might drive tumor progression by releasing hyperactive ERK. We sought to identify the molecular mechanism that upregulates ERK activity during *NKX2-1* repression and the mechanism’s contribution to tumor growth and invasion.

ERK mediates negative feedback signaling to multiple upstream components of the RAS pathway. In this way, cellular ERK activity is highly constrained, even in cells expressing mutant, constitutively active KRAS (31). *In vivo*, release from negative feedback loops allows for elevated ERK signaling that drives malignant progression (32–34). Negative feedbacks in LUAD include ERK’s induction of *Dual-specificity MAPK phosphatases (DUSPs)* and *SPROUTY* proteins *(SPRYs)* (33). The DUSP6 phosphatase specifically dephosphorylates the activation loop of ERK, thereby inactivating ERK (35–37). DUSP6 is upregulated in early lung cancer lesions with activating *EGFR* or *RAS* mutations (38) and is part of a five-gene signature that predicts relapse-free and overall survival in patients with non-small cell lung cancer (NSCLC), of which LUAD is the major subtype (39). *DUSP6* expression is frequently lost in LUAD progression (40), which suggests that reducing DUSP6 levels releases ERK activity for tumor-promoting signaling. Reduced DUSP6 can also promote resistance to ERK pathway inhibitors (41). SPRY proteins inhibit ERK activity by inhibiting RAS and RAF activation (33, 42). Somatic KRAS^G12D^ mutation in mice induces SPRY2 expression (18, 24). *SPRY2* expression is reduced in human LUAD compared to adjacent normal tissue (43). In mice, loss of *Spry2* increases KRAS^G12D^-driven tumor burden (18). Thus, DUSP6 and SPRY2 are candidate RAS/ERK negative feedback loops that must be overcome for progression to LUAD.

KRAS mutations are inversely correlated with *NKX2-1* expression (44, 45), suggesting RAS-driven LUADs undergo selective pressure to lose NKX2-1. We previously found that engineered deletion of *NKX2-1* in KRAS^G12D^-driven LUAD causes a 2-fold reduction in *DUSP6* message (*p*=0.047) and a trend for a 2-fold reduction in *SPRY2* message (*p*=0.103) (24). Further, *NKX2-1* and *DUSP6* expression positively correlate in human LUAD tumor samples and cell lines (46). Thus, we hypothesized that in RAS-driven LUAD, *NKX2-1* silencing is a mechanism to remove *DUSP6* and/or *SPRY2* expression and elevate ERK activity for tumor growth and metastasis.

We found that NKX2-1 loss in LUAD clinical samples and cell lines correlates with reduced DUSP6. Further, NKX2-1 directly induces *DUSP6* expression and limits tumor cell proliferation, migration, invasion, and tumor growth metastasis. While *DUSP6* knockout can be toxic, it abrogates the effects of NKX2-1 on cell proliferation, migration and tumor growth. Inducing DUSP6 in KRAS-driven tumors shows that the DUSP6-feedback loop is sufficient for NKX2-1 mediated tumor suppression. Thus, NKX2-1 tempers ERK activity and lung adenocarcinoma progression through the induction of the ERK phosphatase DUSP6.

## Results

### NKX2-1 transcriptionally induces *DUSP6*

We hypothesized that NKX2-1 promotes the DUSP6 or SPRY2 negative feedback loops that temper ERK signaling in LUAD. In this model, *NKX2-1* suppression would weaken the negative feedback and release high RAS-driven ERK activity. Retention of NKX2-1 would maintain low ERK activation and higher DUSP6 and SPRY2 expression. To test this, we first compared the amounts of *NKX2-1*, *DUSP6*, and *SPRY2* expression in human LUAD using RNA-seq data from The Cancer Genome Atlas (2). We found that *NKX2-1* mRNA levels positively correlate with both *DUSP6* and *SPRY2* mRNA (Fig. S1A, *DUSP6* r^2^=0.28, *p*=1.8 e^−12^, *SPRY2* r^2^=0.33, *p*=1.2 e^−16^). The relatively low r^2^ values indicate that other factors also contribute to *DUSP6* and *SPRY2* expression, which is expected given their regulation by multiple other transcription factors including the ERK-activated ETS and CREB factors (20, 47–49). *DUSP6* and *SPRY2* mRNA levels were highly correlated (Fig. S1B, r^2^=0.49, *p*=1.4 e^−36^), as they are both similarly induced by ERK (33). We next tested if NKX2-1 protein expression correlates with DUSP6 and/or SPRY2 in human samples of LUAD. We stained human tumor microarrays for NKX2-1, DUSP6, and SPRY2 and scored their intensities on a scale of 0 (no staining) to 3+ (highest staining). Samples with NKX2-1 intensities of 2+ and 3+ contained significantly more DUSP6 than samples lacking NKX2-1 (Fig. 1A, *p*=0.001 and 0.002). No association was found between SPRY2 and NKX2-1 (Fig. 1A). Thus, NKX2-1 expression correlates with DUSP6 levels, but not SPRY2 levels.

**Figure 1.**
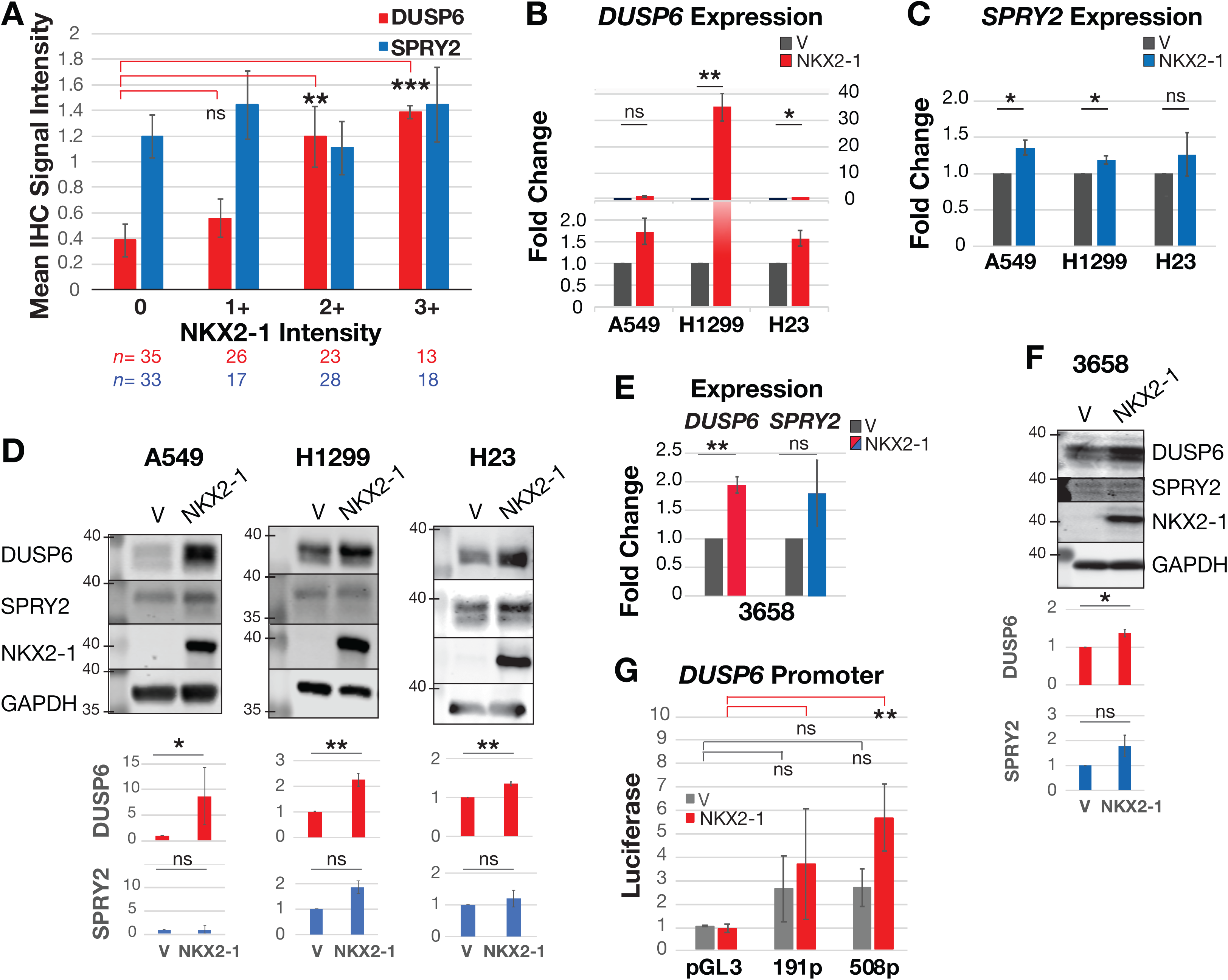
NKX2-1 transcriptionally induces *DUSP6*. **A.** Immunohistochemistry (IHC) intensity scores of DUSP6 and SPRY2 in human LUAD TMAs. Significance calculated with two-sided Dunnett test with 95% CI. **B.** and **C.** qRT-PCR of *DUSP6* and *SPRY2* upon NKX2-1 expression in human RAS mutant, NKX2-1-negative LUAD cell lines: A549 KRAS^G12S^ *n*=3, H1299 NRAS^Q61K^ *n*=3, and H23 KRAS^G12C^ *n*=4 biological replicates. Values are mean and SEM. **D.** Representative Westerns and quantification of DUSP6 and SPRY2 levels upon exogenous NKX2-1 expression in LUAD cell lines. V = empty vector. Relative expression compared to GAPDH. Means and SD from *n*=4 independent experiments for A549, *n*=3 for H1299 and H23. **E.** qRT-PCR of *DUSP6* and *SPRY2* upon NKX2-1 expression in mouse 3658 cells, KRAS^G12D^ *n*=3. Mean and SEM. **F.** Representative Westerns of DUSP6 and SPRY2 levels upon NKX2-1 expression in 3658 cells. Relative expression compared to GAPDH. Means and SD from *n*=3 independent experiments. **G.** *DUSP6* luciferase reporter assay using A549 cells. Mean and SEM of normalized luciferase from *n*=3 independent experiments. Significance of differences in means in RT-PCR, Western, and luciferase reporter data tested with one-way ANOVA and Tukey-Kramer test, 95% CI.

We tested if NKX2-1 is sufficient to induce DUSP6 or SPRY2 mRNA and protein levels in a panel of human LUAD cell lines that contain RAS oncogenes and silenced *NKX2.1*. Quantitative reverse transcription PCR (qRT-PCR) showed that exogenous NKX2-1 expression increased *DUSP6* mRNA (A549 trend with *p*=0.069, H1299 *p*=0.003, H23 *p*=0.020, Fig. 1B). NKX2-1 increased *SPRY2* mRNA in A549 and H1299 cells (*p*=0.026 and 0.034), but not in H23 cells (*p*=0.414, Fig. 1C). Western blotting showed that re-expression of NKX2-1 induced DUSP6 protein levels (A549 *p*=0.033, H1299 *p*=0.007, H23 *p*=0.002, Fig. 1D). NKX2-1 did not induce SPRY2 protein in any cell line (A549 *p*=0.617, H1299 *p*=0.098, H23 *p*=0.493, Fig. 1D).

We also tested if NKX2-1 induces *DUSP6* or *SPRY2* in a murine LUAD cell line derived from KRAS^G12D^; TP53^Null^; NKX2-1^Null^ mouse tumors (3658 cells, (24)). Indeed, qRT-PCR again showed that exogenous NKX2-1 increased *Dusp6* mRNA (*p*=0.003), but not *Spry2* mRNA (*p*=0.238, Fig. 1E). Similarly, NKX2-1 increased DUSP6 protein levels (*p*=0.022), but not SPRY2 (*p*=0.140, Fig. 1F). Together, these data show that NKX2-1 induces DUSP6 consistently across species and LUAD genotypes.

Since NKX2-1 regulates *DUSP6* expression, we sought to determine if NKX2-1 directly controls *DUSP6* transcription. The promoter region of *DUSP6* is highly conserved in vertebrates and located approximately 1000-250 bp upstream of the transcription start site (*Mm*, (48)). Our previous chromatin immunoprecipitation-sequencing (ChIP-seq) of NKX2-1 in KRAS^G12D^ mouse tumors showed NKX2-1 binds within the promoter region of *DUSP6*, with the greatest enrichment between −600 and −400 (Fig. S1C, (24)). ChIP-seq of NKX2-1 in human LUAD lines showed NKX2-1 binds promoters in areas proximal to AP-1 and Forkhead box (FOX) binding motifs (27, 46). We assayed in A549 cells the luciferase reporter activity of a control promoter region that included the putative transcription start site at −463 and a portion of the putative NKX2-1 binding region (191p, −550 to −359) and larger region that included the entire binding region for NKX2-1 (508p, −866 to −359). In the presence of NKX2-1, the 508p region exhibited 5 times more transcriptional activity than the pGL3 vector (*p*=0.011, Fig. 1G). NKX2-1 did not induce the activity of the shorter 191p region (*p*=0.275). Thus, in human lung adenocarcinoma, NKX2-1 appears to directly induce *DUSP6* gene expression and increase DUSP6 protein levels.

### NKX2-1 inhibits cell proliferation, migration, and invasion

If NKX2-1 inhibits tumor progression in part by inducing DUSP6 to suppress ERK activity, then NKX2-1 should inhibit ERK-mediated and DUSP6-regulated cancer phenotypes, such as cell proliferation, migration, and invasion, and tumor growth and dissemination. To test this hypothesis, we assayed the proliferation of our *NKX2-1-*silenced LUAD cells complemented with NKX2-1. NKX2-1 expression uniformly inhibited cell proliferation in A549, H1299, H23, and 3658 cells (Fig. 2A, S2A).

**Figure 2.**
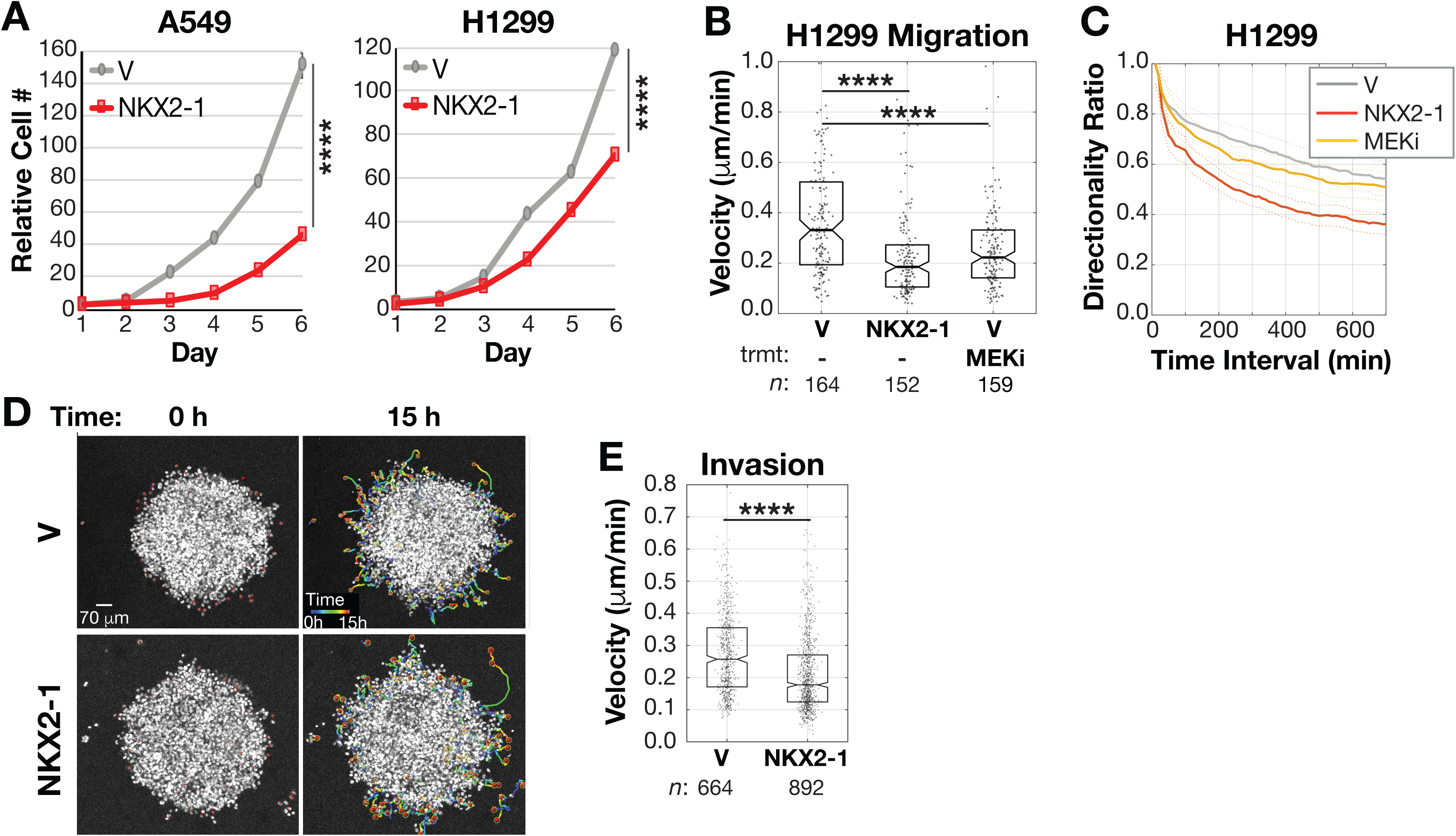
NKX2-1 inhibits cell proliferation, migration, and invasion. **A.** Proliferation of A549 and H1299 cells. Significance from one-way ANOVA. **B. and C.** Random-walk migration velocity and directionality of H1299 cells upon NKX2-1 expression and MEK inhibition (Selumetinib 10 μM). Velocity is Mean Squared Displacement. Significance from two-sample Kolmogorov-Smirnov (K-S) test. **** indicates *p*<0.0001. **D.** Representative images from invasion assay. Central mass is spheroid and colored lines are tracks of invading cells. **E.** Invasion velocity as distance/time from 4 independent experiments. Significance from K-S test.

We sought to test if NKX2-1 also inhibits cell migration and invasion. We previously found that inhibition of MEK (and therefore ERK) reduces cell migration and persistence in multiple cell lines, including A549 and 3658 cells (50). However, A549 and 3658 cells are poorly migratory compared to other assayed lines (untransformed Cos7 and MDCK, and fibrosarcoma HT1080 cells (50)). Using a random walk migration assay and automated tracking, we found that of the four KRAS mutant, *NKX2-1* silenced LUAD cell lines, H1299 cells exhibit the fastest 2D motion, with MSD velocity of 0.24 μm/min (Fig. S2B). We therefore used H1299 cells to test if NKX2-1 controls cell migration. NKX2-1 re-expression in H1299 cells slowed migration velocity from 0.33 μm/min in cells with empty vector to 0.19 μm/min in cells expressing NKX2-1 (*p*=1.8×10^−9^, Fig. 2B). Consistent with our prediction that NKX2-1 expression would phenocopy ERK inhibition, NKX2-1’s inhibition of migration velocity was similar to treatment with the MEK inhibitor Selumetinib, which reduced movement to 0.22 μm/min (*p*=7.1×10^−6^, Fig. 2B). We calculated the persistence, or straightness of the trajectory, as a directionality ratio at each time point. A higher score indicates more linear, productive motion. We found that NKX2-1 also reduces migration directionality (Fig. 2C). We tested if NKX2-1 inhibits invasion in soft, 3D collagen matrices. We embedded H1299 cells as spheroids in collagen I and tracked invasion velocity as the cells moved out into the gel. We found that NKX2-1 re-expression slowed invasion velocity from 0.257 μm/min to 0.177 μm/min (*p*=4×10^−19^, Fig. 2D, E). Thus, NKX2-1 inhibits migration on hard 2D surfaces and invasion through a physiological 3D environment.

We rationalized that if NKX2-1 induces DUSP6, then NKX2-1 expression would reduce the amount of phospho- (p-) ERK. However, previous work in LUAD cell lines with mutant *EGFR* or *KRAS* has shown that manipulation of DUSP6 has a more nuanced effect on p-ERK. After *DUSP6* knockdown, cells in culture show an initial, slight increase in p-ERK levels (1.5-fold within 24 hr), as expected from removing the negative regulator (38). However, the regulation is transient. After 5 days of *DUSP6* knockdown, cells in culture show decreased p-ERK and toxicity (38). In normal human bronchial epithelial cells (NHBE), KRAS^V12^ expression suppresses growth, but surviving subpopulations with increased p-ERK emerge (40). When combined with dominant negative DUSP6 mutant C293S, the surviving subpopulations appear to harbor even more p-ERK (40). Thus, cells with RAS mutations adjust their feedback signaling to constrain p-ERK, and long-term selective pressure can lead to genetic or epigenetic changes that allow cells to benefit from reduced DUSP6 activity and increased p-ERK (31, 38, 40). In our LUAD cells with re-introduction of NKX2-1, we found that the p-ERK induced by stimulation with RAS pathway agonists EGF or PMA trended lower in cells expressing NKX2-1 compared to cells with empty vector (Fig. S2C, S2D, not significant). The small change is consistent with the previous studies on DUSP6.

### NKX2-1 induces DUSP6 and inhibits p-ERK during tumor progression

We further tested if NKX2-1’s inhibition of cancer phenotypes is associated with *DUSP6* induction and ERK inhibition *in vivo*, in which more stringent biological pressures for growth and survival model the pathway rewiring of tumorigenesis. Our and others’ previous studies in transgenic mice showed that NKX2-1 limits KRAS^G12D^-driven LUAD progression (24, 25, 27). We also showed that *Nkx2-1* deletion in KRAS-driven tumors reduces *Spry2* expresion and induces ERK activation (24). This experiment used *Kras^LA2^; RosaCre^ERT2^; Nkx2-1^F/F^* mice, in which tumors arise via spontaneous recombination of a latent *KRas^G12D^* allele and tamoxifen injection deletes *Nkx2-1* (5, 24). Since our qRT-PCR and Western data suggested that DUSP6 may be the more significant ERK feedback effector in LUAD, we tested if DUSP6 is lost along with NKX2-1. We used *KRas^frtSfrt-G12D/+^; Nkx2-1^F/F^; Rosa^frtSfrt-CreERT2^* mice and intratracheal delivery of FlpO recombinase to activate the *KRas^G12D^* oncogene and express *Cre^ERT2^* Cre recombinase. Following 1 week of tumor initiation, mice were treated with Tamoxifen to activate Cre and delete *Nkx2-1*. After 20 weeks of tumor initiation, lungs were harvested and NKX2-1 positive and negative tumors were identified by immunohistochemistry (IHC) staining for NKX2-1 and its target pro-surfactant protein B (pro-SPB, Fig. S3A). We found that indeed, NKX2-1 deletion reduces DUSP6 along with reducing SPRY2 (Fig. S3A). In agreement with previous results, NKX2-1 deletion increased p-ERK and downstream effectors p-RSK and p-S6 and caused tumors to transition to invasive mucinous adenocarcinoma (Fig. S3B). Thus, NKX2-1 is necessary for the expression of *DUSP6* and suppression of ERK during LUAD tumorigenesis. We note that NKX2-1’s regulation of SPRY2 in the transgenic model contrasts with the LUAD cell lines.

We next tested if NKX2-1 is sufficient for inhibition of tumor growth and metastasis, induction of *DUSP6*, and inhibition of ERK. We generated subcutaneous tumors with the A549 cell line pairs and assessed primary tumor size and frequency of metastasis to the lung. Tumors expressing NKX2-1 were significantly smaller by weight than tumors lacking NKX2-1 (A549-V median tumor weight 0.64 g versus A549-NKX2-1 median 0.17 g, *p*=0.02, Fig. 3A). When we harvested the lungs and manually examined sections for micrometastases, we found that NKX2-1 reduces the occurrence of metastasis. 5 out of 9 mice with A549-V tumors exhibited micrometastases versus 0 out of 9 mice with A549-NKX2-1 tumors (Fig. 3B). IHC showed that tumors lacking NKX2-1 exhibit low DUSP6 expression and high p-ERK (Fig. 3C). In contrast, tumors with NKX2-1 reconstitution exhibited high *DUSP6* expression and reduced p-ERK (Fig. 3C). SPRY2 was not regulated (Fig. 3C).

**Figure 3.**
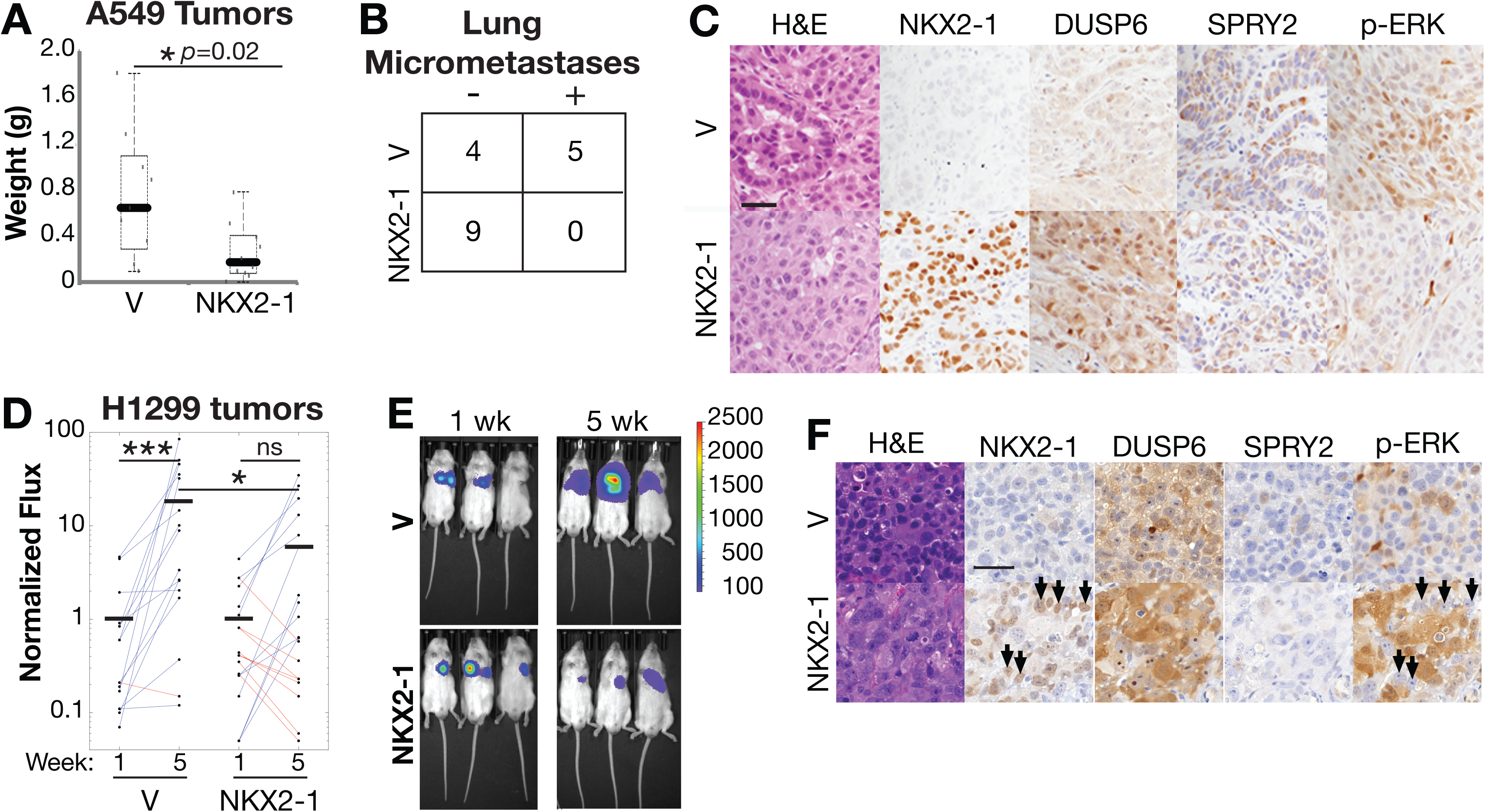
NKX2-1 limits ERK activity and tumor growth and dissemination. **A.** Tumor weight from A549 cells transplanted subcutaneously in mice. *p*=0.02. Central line is median. Lower and upper box limits are 1^st^ and 3^rd^ quartiles. Dashed bars span minimum and maximum. Significance by Wilcox Mann-Whitney test. **B.** Number of mice (9 total per condition of V or NKX2-1) with lung A549 micrometastases. Fisher’s exact test *p*=0.0294, Chi-square *p*=0.0085. **C.** Representative images of hematoxylin and eosin (H&E) staining to show cells and extracellular matrix, and IHC staining of NKX2-1, DUSP6, SPRY2, and p-ERK in primary A549 tumors. Scale bar 100 μm. **D.** Tumor size in H1299 orthotopic transplants, indicated by total luciferase flux and normalized to initial size at week 1. Blue tumors grew. Red tumors shrank. Horizontal black lines show median. *n*=16 mice with V, 19 mice NKX2-1. Significance by Wilcox Mann-Whitney test of medians. **E.** Representative H1299 IVIS. **F.** IHC of stable H1299 cells transplanted orthotopically into mouse lungs. Representative images of H&E, NKX2-1, DUSP6, SPRY2, and p-ERK stains. Scale bar 50 μm.

We also tested the growth of H1299 cells transplanted orthotopically into mouse lungs. Our qRT-PCR and Western data showed that H1299 cells exhibit higher basal expression of DUSP6 (Fig. 1). The cell line pairs with empty vector and NXK2-1 were infected for stable expression of GFP-Luciferase, injected into the lung, and followed by bioluminescence imaging. After 5 weeks of growth, we found that tumors without NKX2-1 grew 18-fold (H1299-V normalized median flux 1 photons/sec increased to 18.5 photons/sec, *p*=0.001, Fig. 3D, 3E). In contrast, tumors expressing NKX2-1 did not exhibit significant growth (H1299-NKX2-1 normalized median flux at 1 week versus 5 weeks, *p*=0.28, Fig. 3D, 3E). Rather, after 5 weeks H1299-NKX2-1 tumors were 3 times smaller than H1299-V tumors (H1299-NKX2-1 average 6.1 photons/sec, *p*=0.04, Fig. 3D, 3E). IHC showed a more complicated scenario of subpopulations with moderate and high NKX2-1 expression (Fig. 3F). Cells with moderate NKX2-1 harbored high DUSP6 expression, but high p-ERK. In contrast, cells with high NKX2-1 harbored moderate increases in DUSP6 and no p-ERK (Fig. 3F, arrows). SPRY2 levels were unchanged. We surmise that the unexpected population with moderate NKX2-1, but high DUSP6 and p-ERK, is a resut of re-wiring that strengthened the ERK→ETS→DUSP6 signal (20) relative to the NKX2-1→DUSP6 signal. In sum, these A549 and H1299 transplant data are consistent with our hypothesis that NKX2-1-mediated DUSP6 induction limits ERK activation to temper ERK-mediated tumor progression.

### NKX2-1 requires DUSP6 to inhibit cell proliferation and migration

We sought to directly test if NKX2-1 acts through DUSP6 or SPRY2 to control tumor cell proliferation and migration. Previous work knocking down DUSP6 in NHBE and A549 cells did not show regulation of cell growth, suggesting that knockdown efficiency was insufficient to overcome KRAS-induced activation of ERK and expression of DUSP6 (40). Therefore, we generated A549 and H1299 *DUSP6* and *SPRY2* CRISPR/Cas9 knockouts, confirmed by Western (Fig. S4A, S4C). While DUSP6’s role as a negative feedback regulator of ERK suggests that DUSP6 loss should result in faster cell proliferation and migration, complete loss of DUSP6 can be toxic in LUAD cells that harbor KRAS and EGFR mutations (38). We found that in A549 cells, *DUSP6* and *SPRY2* CRISPR/Cas9 knockout appeared to slow A549 cell proliferation by Day 5, although the trend was not significant (Fig. S4B). In H1299 cells, *DUSP6* loss did not change cell proliferation (Day 6 *p*=0.38), but *SPRY2* loss increased proliferation (Day 5 *p*=7.5×10^−6^, Fig. S4D). This suggests that in cell culture, A549 and H1299 cells engage compensatory signaling to limit the effects of DUSP6 loss. However, under these unpressured growth conditions, SPRY2 tempers cell proliferation.

We tested if NKX2-1 requires DUSP6 or SPRY2 to inhibit proliferation by introducing TRE-NKX2-1 into *DUSP6* and *SPRY2* knockout clones of A549 and H1299 cells. Doxycycline induced *NKX2-1* expression and *DUSP6* expression (Fig. 4A, B). As before, p-ERK was not significantly changed *in vitro* (Fig. S5A, B). In A549 cells, NKX2-1 expression slowed the proliferation of the control CRISPR cells 2.4-fold (Day 6, *p*=0.01), but not the already slow *DUSP6* or *SPRY2* knockouts (Fig. 4C). Similarly, in H1299 cells, NKX2-1 expression slowed the proliferation of the control GFP CRISPR cells 1.6 fold (Day 6, *p*=0.03), but not the *DUSP6* or *SPRY2* knockouts (Fig. 4D). We conclude that NKX2-1 requires both DUSP6 and SPRY2 to fully suppress proliferation *in vitro*.

**Figure 4.**
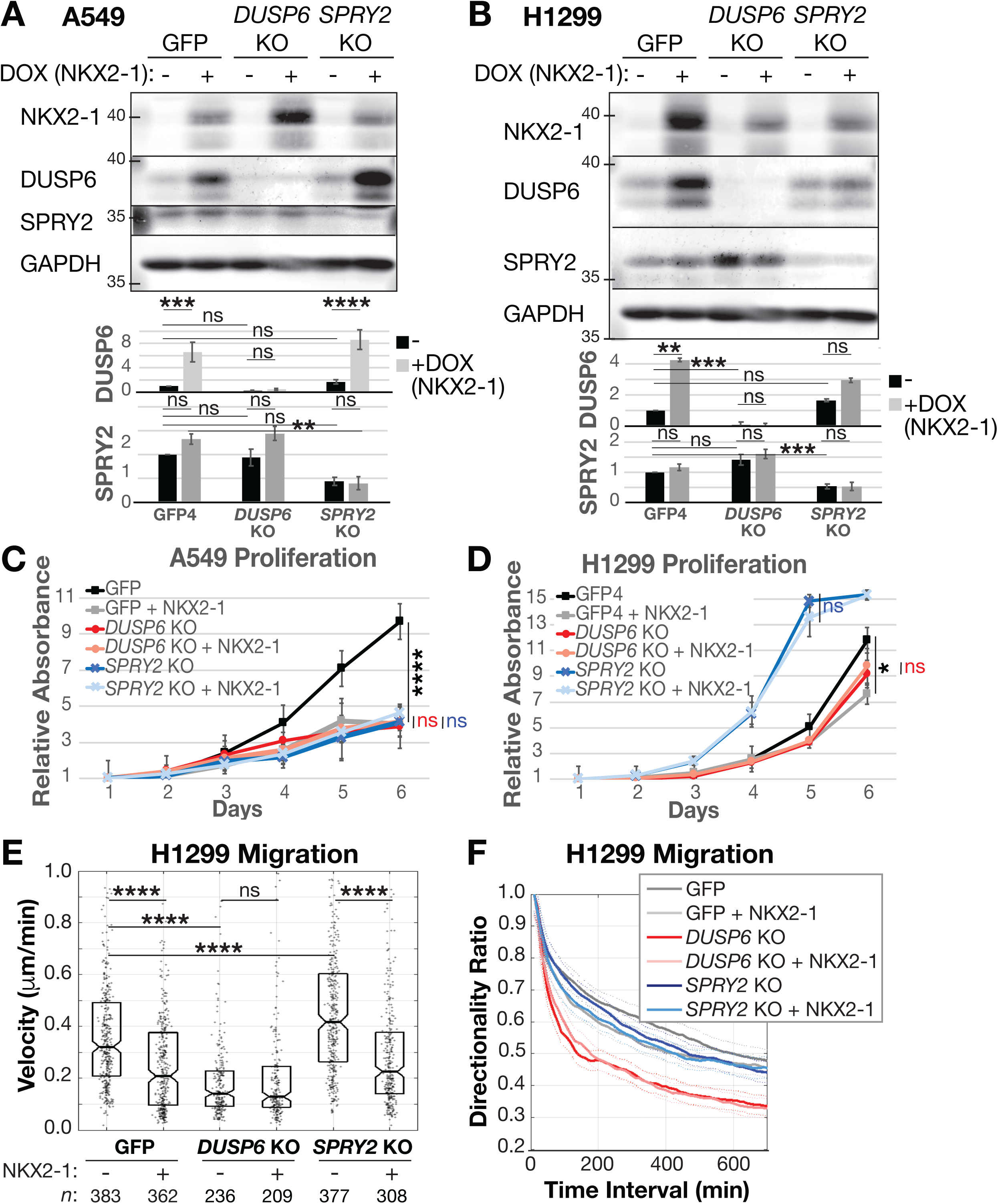
NKX2-1 requires DUSP6 to slow cell proliferation and migration. **A.** and **B.** Westerns and quantification of A549 and H1299 *DUSP6* and *SPRY2* knockouts (KOs) with doxycycline (DOX) induction of TRE-NKX2-1. Relative expression compared to GAPDH. Means and SD from *n*=3 independent experiments. **C.** and **D.** Cell proliferation upon NKX2-1 induction in A549 and H1299 *DUSP6* and *SPRY2* KOs. Significance between uninduced and DOX-induced NKX2-1 expression for each cell line calculated by one-way ANOVA. GFP and *DUSP6* KO significance at day 6. *SPRY2* KO significance at day 5. *n*=3 experiments. **E.** and **F.** H1299 cell migration velocity and directionality upon NKX2-1 induction. Significance from two-sample Kolmogorov-Smirnov (K-S) test, **** indicates *p*<0.0001.

We tested if NKX2-1 regulates cell migration using the H1299 cells. *DUSP6* knockout slowed cell migration velocity (control CRISPR targeting GFP 0.34 μm/min versus *DUSP6* knockout 0.19 μm/min, *p*=7.4×10^−16^) and reduced migration directionality (Fig. S4E, S4F). We tested additional *DUSP6* knockout clones and found they uniformly exhibited slower migration velocity and reduced directionality (Fig. S4G, S4H). In contrast, *SPRY2* knockout increased migration velocity (0.52 μm/min, *p*=1.1×10^−19^, Fig. S4E). We then tested if NKX2-1 suppressed migration in the absence of DUSP6 or SPRY2. Doxycycline-mediated NKX2-1 expression slowed the migration of control CRISPR cells from 0.32 μm/min to 0.21 μm/min (*p*=1.5×10^−13^, Fig. 4E). NKX2-1 also slowed the migration of the *SPRY2* knockouts (0.42 μm/min slowed to 0.23 μm/min, *p*=8.5×10^−17^, Fig. 4E). However, NKX2-1 expression did not further slow the migration of the *DUSP6* knockouts. *DUSP6* knockout migration of 0.14 μm/min was unchanged with NKX2-1, *p*=0.67, Fig. 4E). Regulation of migration directionality followed a similar pattern (Fig. 4F). Thus, NKX2-1 requires DUSP6, but not SPRY2, to inhibit cell migration.

### NKX2-1 drives tumor progression through DUSP6

While NKX2-1 required both DUSP6 and SPRY2 to control *in vitro* cell proliferation (Fig. 4C, 4D), NKX2-1 primarily induced *DUSP6* expression in human samples and cell lines (Fig. 1) and only DUSP6 was required for NKX2-1 suppression of migration (Fig. 4E). Thus, we hypothesized that NKX2-1 acts through DUSP6 to promote LUAD progression *in vivo*. To test this, we generated subcutaneous tumors with the A549 *DUSP6* knockouts harboring inducible TRE-NKX2-1 and GFP-luciferase. After the tumors reached 150-400 mm^3^ (5 weeks for control tumors and 7 weeks for DUSP6 knockout tumors that initially grew more slowly), we induced NKX2-1 expression with doxycycline. In the week before induction, control tumors grew 8.5-fold (A549-GFP normalized median flux 1 photons/sec increased to 8.5 photons *p*=0.008) while DUSP6 knockout tumors did not appreciably grow (Fig. 5A, B). After 3.5 weeks of NKX2-1 induction, control tumors had stopped growing, with final flux 3.3 photons. In contrast, DUSP6 knockout tumors exhibited the same slow growth trend (Fig. 5A, B). This indicates that NKX2-1 inhibition of tumor growth requires DUSP6.

**Figure 5.**
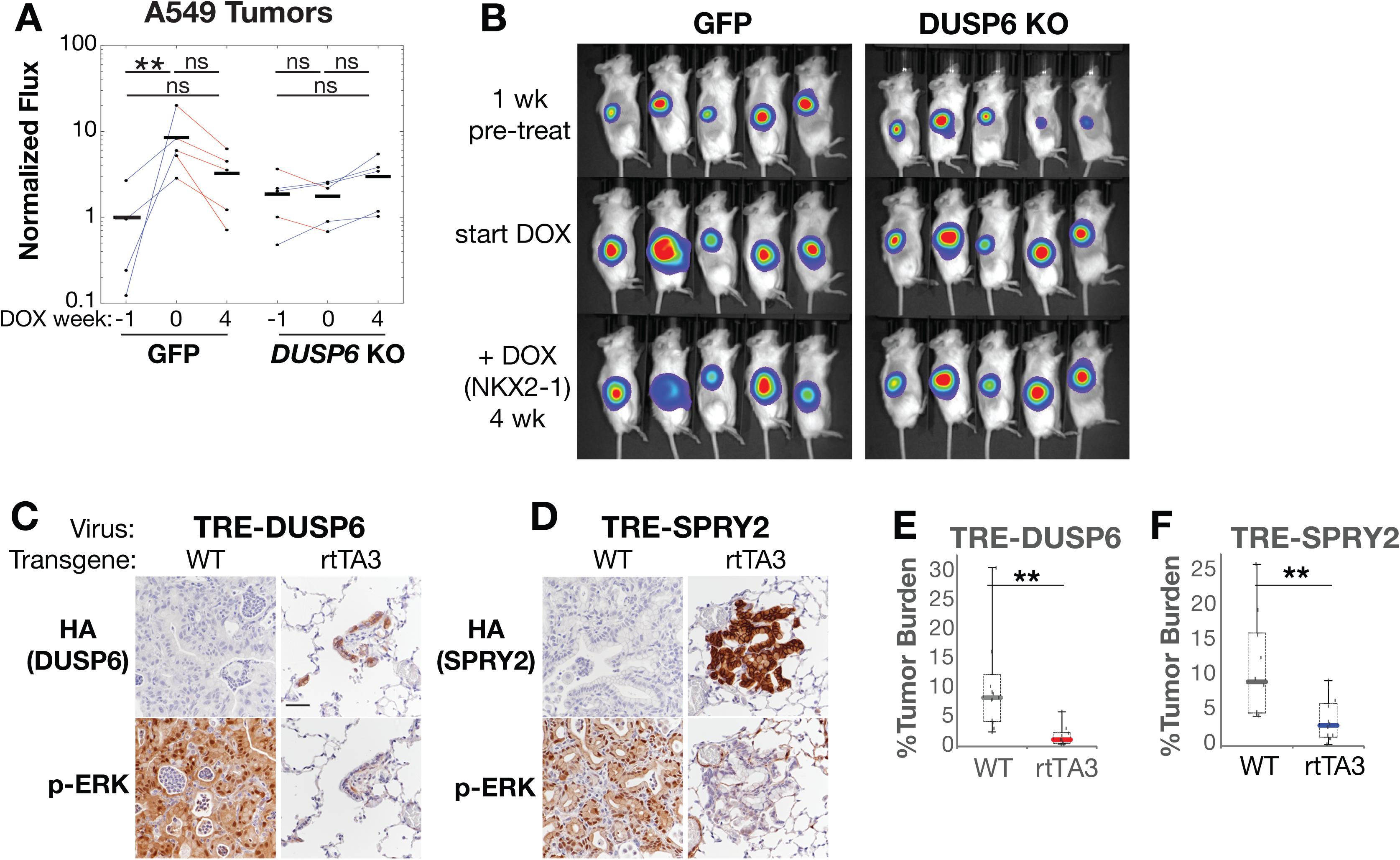
NKX2-1 controls tumor progression through DUSP6. **A.** Tumor size in A549 subcutaneous tumors, indicated by total luciferase flux and normalized to initial size before induction of NKX2-1 with DOX. Horizontal black lines show median flux at 1 week before induction (week -1), at induction (week 0), and 4 weeks after induction (week 4). *n*=5 mice with control A549 cells with *GFP* CRISPR and 5 mice with A549 with *DUSP6* knockout. Significance by Wilcox Mann-Whitney test of medians. **B.** IVIS images for time point in A. **C.** and **D.** IHC for p-ERK in KN-rtTA3 mice infected with TRE-Dusp6 and TRE-Spry2, after 1 week of doxycycline-mediated de-repression of rtTA3 and induction of DUSP6 or SPRY2. **E.** and **F.** Tumor burden in mice with KRAS^G12D^;NKX2-1^Null^; WT tumors and KRAS^G12D^;NKX2-1^Null^; rtTA3 treated with TRE-Dusp6 and TRE-Spry2 for 1 week.

Lastly, we tested if DUSP6 or SPRY2 overexpression is sufficient to temper LUAD tumor growth. We generated tumors in *Kras^LSL-G12D/+^; Nkx2-1^F/F^* control mice and *Kras^LSL-G12D/+^; Nkx2-1^F/F^ CAG-rtTA3* mice (8, 51) using a dual promoter lentivirus that encodes constitutive Cre and doxycycline-inducible *Dusp6* or *Spry2*. The *CAG-rtTA3* transgene drives ubiquitous expression of the tetracycline-regulated transactivator gene. After 5 weeks of tumor growth, we used a doxycycline diet to induce exogenous DUSP6 or SPRY2 in established KRAS^G12D^, NKX2-1^Null^ tumors. After 1 week of treatment, we found that in control mice lacking *rtTA3*, the tumors resembled those of our *KRas^frtSfrt-G12D/+^;Trp53^frt/frt^;Nkx2-1^F/F^* mice: low DUSP6 expression and high, uniform p-ERK (Fig. 5C wildtype, Fig. S3A, S3B NKX2-1^Null^). The tumors in mice with the *rtTA* transgene showed heterogeneous HA-tagged DUSP6 and SPRY2 expression, likely due to stochastic silencing of integrated lentivirus following tumor initiation (Fig. 5C, D). Individual tumors that successfully induced DUSP6 with one week of doxycycline chow had regressed to small lesions with low p-ERK that resemble the alveolar hyperplasia induced by *Nkx2-1* deletion alone (24). Interestingly, induction of DUSP6 caused some decrease in SPRY2, and induction of SPRY2 caused some decrease in DUSP6 levels (Fig. S5C, D). This suggests that *in vivo* pressures are actively constraining p-ERK. We quantified tumor burden in the lung by histological analysis and found that DUSP6 expression reduced overall tumor burden (Fig. 5E). Consistent with our previous finding that NKX2-1 deletion reduces SPRY2 (24) in mouse tumors, SPRY2 expression reduced tumor size and p-ERK in this system (Fig. 5F). In summary, DUSP6 and SPRY2 are sufficient to limit tumorigenesis in the genetically engineered model of LUAD.

## Discussion

Activating mutations in *EGFR*, *KRAS* and *BRAF* initiate LUAD tumors by enhancing ERK signaling to drive cell survival, proliferation, and migration (52). Mouse models of LUAD suggest that an ERK pathway negative feedback must be weakened for early KRAS^G12D^ lesions to acquire tumor-promoting ERK activity (5, 8). Discerning the mechanisms of negative feedback disruption is a high priority, as they could be targeted to prevent tumor progression. One mechanism likely involves *KRAS* allelic imbalance, in which mutant *KRAS* is amplified and the wildtype *KRAS* allele is inactivated (12, 17). Allelic imbalance has been documented in the KRAS^G12D^; p53^Null^ tumors in mice, in some human cancer cell lines (12, 53), and in 2% of human LUADs (19, 54). However, how ERK activity is increased to levels that promote tumor progression in the remaining LUADs and mechanisms specific to LUAD histopathologies and differentiation states are unknown.

Here, we identify a mechanism of ERK feedback disruption specific to *KRAS*-driven mucinous LUAD transitioning through dedifferentiation. NKX2-1 is expressed in alveolar type II cells in the adult, the predominant cell of origin for LUAD (55–57). We show that NKX2-1 directly induces DUSP6. 5-10% of LUADs are diagnosed as invasive mucinous (21) and loss of NKX2-1 leads to invasive mucinous adenocarcinoma with pulmonary to gastric transdifferentiation (24). We show in a genetically-engineered mouse model of LUAD that DUSP6 is downregulated concomitant with NKX2-1. In LUAD cell lines and xenografts, NKX2-1 re-expression causes DUSP6 upregulation. Further, DUSP6 is both necessary and sufficient for NKX2-1 to inhibit *in vivo* ERK activity and tumor growth. These findings lead to the conclusion that in early KRAS mutant lesions, NKX2-1’s induction of DUSP6 maintains the negative feedback signaling of the RAS→RAF→MEK→ERK pathway and thereby limits ERK activity and tumor progression. NKX2-1 silencing and reduced DUSP6 levels would allow for increased ERK activity that promotes tumor progrssion during the transdifferentiation process.

In addition to DUSP6, SPRY2 also appears to play a role in ERK pathway feedback in mouse LUAD. Previous analysis of human tumors in the TCGA found that *DUSP6* was the only ERK pathway negative-feedback regulatory gene expressed differently in tumors with *KRAS* or *EGFR* mutations versus tumors without (38). Consistent with this human study, we found that DUSP6, but not SPRY2 levels correlated with NKX2-1 in human LUAD samples. Further, DUSP6 but not SPRY2, protein was significantly upregulated upon introduction of NKX2-1 into human LUAD cell lines, both *in vitro* and in xenografts. However, in the genetically engineered mouse tumors, *Nkx2-1* deletion caused a loss of SPRY2 along with a loss of DUSP6. This suggests that NKX2-1 works through DUSP6 in the human disease, but in the KRAS^G12D^ mouse model NKX2-1 may utilize both DUSP6 and SPRY2 to suppress tumor progression. Regardless of regulation by NKX2-1 or not, SPRY2 contributes to the homeostatic regulation of proliferation.

Loss of *DUSP6* expression likely occurs more broadly across the LUAD histopathologies. In general, lower DUSP6 expression correlates with higher tumor grade and reduced overall survival (58). In addition to being a transcriptional target of the ERK-induced ETS transcription factors, and NKX2-1 shown here, *DUSP6* is a target of p53 (59, 60). p53 loss occurs in 20-40% of LUADs, except the recently defined proximal-proliferative subcluster (2, 44, 61). Thus, it will be important for future studies to determine if p53 silencing participates in removing the DUSP6 negative feedback loop in non-mucinous LUADs.

Different cell types in the lung have distinct thresholds for oncogenic versus toxic ERK activity, suggesting that NKX2-1-regulated DUSP6 could contribute to a cell type-specific transformation process. Alveolar type II cells transformed by dual *KRas* and *BRaf* mutations suffer toxicity from excessively high p-ERK that occurs with loss of the wildtype *BRaf* allele (16). In contrast, club cells are transformed by the same high p-ERK and develop into intrabronchiolar lesions (16). Our data suggest that in KRAS-transformed alveolar type II cells, silencing of NKX2-1 and the resulting reduction in DUSP6 levels increases p-ERK to a sweet spot of tumor-promoting activity without toxic hyperactivity. However, complete knockout of DUSP6 *in vitro* had the opposite effect: cell proliferation was unchanged and cell migration was slowed. These latter data are consistent with previous *in vitro* findings that nearly complete knockdown of DUSP6 in cells with *RAS* or *EGFR* mutations induces cell toxicity (38, 62, 63). Surprisingly, the same cells showed tumor growth *in vivo*, suggesting that under selective pressure, the cells rewire to adopt a p-ERK signal intensity that promotes growth and dissemination.

NKX2-1’s regulation of DUSP6 to control ERK activation has therapeutics implications. We recently found that BRAF^V600E^; NKX2-1^WT^ tumor cells exit the cell cycle when treated with BRAF and MEK inhibitors, but NKX2-1-negative persister cells arrest within the cell cycle (64). Together, these data suggest that the NKX2-1-positive cells are addicted to the lower level of ERK activity maintained in the presence of NKX2-1-induced DUSP6. Similarly, LUAD cells with acquired resistance to EGFR inhibitors are addicted to the inhibitor-dampened ERK activity such that inhibitor withdrawal induces toxic ERK hyperactivation (65). Lineage heterogeneity promotes chemoresistance (26). Models of tumors with heterogeneous ERK activity show that ERK heterogeneity contributes to differences in transcriptional states that promotes tumorigenesis thorugh paracrine signaling (66). This suggests the acquisition of an NKX2-1-negative subpopulation would contribute to both lineage and ERK activity heterogeneity, both of which would complicate treatment. Therapeutic strategies to inhibit ERK activity need to attain a uniform reduction in p-ERK, and upfront inhibition of MEK along with the upstream oncogene can forestall resistance (41). Alternative strategies to inhibit DUSP6 and induce toxic ERK hyperactivation are also under investigation (67, 68), but due to the heterogeneity and adaptability of the ERK pathway system, will likely be most beneficial as agents that sensitize cells to chemotherapeutics rather than stand-alone treatments.

---

**Figure S1. *NKX2-1* expression correlates with *DUSP6* and *SPRY2* in human LUAD. A. and B.** Correlation analyses of *NKX2-1*, *DUSP6*, and *SPRY2* expression (TCGA LUAD). R value is Pearson’s correlation coefficient. **C.** Representattive signal from ChIP-Seq data for NKX2-1 at *DUSP6* locus (MACS, (69)).

**Figure S2. NKX2-1 inhibits cell proliferation, but p-ERK levels are normalized. A.** Proliferation of H23 and 3658 cells with exogenous NKX2-1. Error bars are SEM. Significance one-way ANOVA, **** indicates *p*<0.0001. **B.** Random-walk migration velocity of LUAD cell lines (A549 *n*=836, H1299 *n*=770, H23 *n*=515, 3658 *n*=916). Significance from two-sample Kolmogorov-Smirnov (K-S) test, **** indicates *p*<0.0001. **C.** and **D.** Westerns and quantification of active, phosphorylated (p-) ERK in LUAD cell lysates with exogenous NKX2-1. Relative expression compared to total ERK. V = empty vector. Error bars are SD. *n*=3 for each cell line. Cells starved for 24 h. EGF 50 ng/ml stimulation for 10 min.

**Figure S3. NKX2-1 is required for DUSP6 expression and ERK activity in vivo. A**. and **B.** Representative IHC images of KRAS^G12D^;NKX2-1^WT^ and KRAS^G12D^;NKX2-1^Null^ tumors: NKX2-1 target pro-SPB, DUSP6, SPRY2, p-ERK and effector p-RSK and p-S6 levels. Scale bars: 100 μm. n=6 mice tamoxifen, n=6 mice corn oil injections.

**Figure S4. DUSP6 and SPRY2 control LUAD cell proliferation and migration. A.** Westerns and quantification of A549 *DUSP6* and *SPRY2* knockouts (KOs). **B.** Proliferation of A549 *DUSP6* and *SPRY2* KOs. Error bars are SEM. Significance one-way ANOVA, *n*=3 experiments. **C.** Westerns and quantification of H1299 *DUSP6* and *SPRY2* KOs. **D.** Proliferation of H1299 *DUSP6* and *SPRY2* KOs. Error bars are SEM. Significance one-way ANOVA, **** indicates *p*<0.0001. *n*=3 experiments. **E.** and **F.** Random-walk migration velocity and directionality of H1299 *DUSP6* and *SPRY2* KOs. Significance from two-sample Kolmogorov-Smirnov (K-S) test, **** indicates *p*<0.0001. **G.** and **H.** Random-walk migration velocity and directionality of three H1299 *DUSP6* KO clones.

**Figure S5. A.** and **B.** Westerns and quantification of p-ERK in A549 and H1299 *DUSP6* and *SPRY2* KOs with DOX induction of TRE-NKX2-1. Relative expression compared to total ERK. Means and SD from *n*=3 independent experiments. **C.** IHC for DUSP6 in TRE-Spry2 tumors and SPRY2 in TRE-DUSP6 tumors.

## Methods

### Histology and IHC

Human tissue microarrays were formalin fixed and paraffin embedded: US Biomax BCS04017, BCS04017a, LC10014a. Arrays were stained for NKX2-1, DUSP6, and SPRY2 using standard methods. Mouse lung lobes were fixed in 10% neutral buffered formalin, processed through 70% ethanol, and paraffin embedded. Sectioning at 4 μm and slide processing, and H&E staining was carried out by the HCI Research Histology Shared Resource. IHC was performed manually using the Vectastain ABC Kit (Vector Laboratories) and Sequenza slide staining racks (Electron Microscopy Sciences). Sections were treated with Bloxall (Vector labs) followed by horse serum (Vector Labs, Burlingame, CA) or Rodent Block M (Biocare Medical, Pacheco, CA), primary antibody, and HRP-polymer-conjugated secondary antibody (anti-rabbit, goat and rat from Vector Labs; anti-mouse from Biocare. The slides were developed with Impact DAB (Vector) and counterstained with hematoxylin. The following primary antibodies were used: NKX2-1 (EP1584Y, Abcam), DUSP6 (EPR129Y, Abcam), SPRY2 (EPR4318(2)(B), Abcam), pERK1/2 (D13.14.4E, CST), pRSK S380 (D3H11), pS6 S235/236 (D57.2.2E, CST), pS6 S240/244 (D68F8, CST), and GFP (D5.1, CST). Images were acquired on a Nikon Eclipse Ni-U microscope with a DS-Ri2 camera and NIS-Elements software or a 3D Histech Pannoramic Midi slide scanner with Case Viewer Software. Tumor quantitation was performed on H&E-stained or IHC-stained slides using NIS-Elements software. All histopathologic analysis was performed by a board-certified anatomic pathologist (E.L.S.). Total tumor burden (tumor area/total area × 100%) and individual tumor sizes were calculated using ImageJ.

### Plasmids

*SPRY2* CRISPR oligonucleotides used in H1299 cells (gRNA target site: GTACTCATTGGTGTTTCGGA) were cloned into pSpCas9(BB)-2A-Puro (Addgene plasmid #62988, cloned by UofU HSC Mutation Generation and Detection Core). *SPRY2* CRISPR oligonucleotides used in A549 cells (gRNA target site: CGTACTGCTCCGCGACCCTG) were cloned into lentiCRISPR v2 plasmid (Addgene pasmid #52961). *DUSP6* CRISPR oligonucleotides used in H1299 cells were cloned into lentiCRISPR v2 plasmid (gRNA target site: GGTATACATTCTGGTTG-GAA).

pBabe H2B-mCherry-Ftractin-eGFP was cloned as follows: pBabePuro-Ftractin-eGFP was first generated by amplifying Ftractin-eGFP from of Ftractin-C1-eGFP (Addgene #58473) using the following primers: (For): AGGCGCGCCTGCCACCATGGCGCGACCACGG and (Rev): GGAATTCCTTATTGTACAGCTCGTCCATGCCG. Amplified fragments were purified, digested with AscI and EcoRI, and ligated into a modified pBabe-Puro retroviral backbone (Addgene plasmid #1764 modified to insert FseI and AscI between BamHI and EcoRI within the MCS, using the following primers (For): GATCCGGCCGGCCCCGCGGATCGATGGCGCGCCG and (Rev): AATTCGGCGCGCCAT-CGATCCGCGGGGCCGGCCG, PMID: 21419341). Second, H2B-mcherry was amplified from H2B mCherry-N2 (PMID 21098116) using the following primers: (For): CCCAAGCTTGGGATGCCAGAGCCAGCGAAG and (Rev): CTAGCTAGCTAGCTTGT-ACAGCTCGTCCATGCC. PCR products were purified and digested with HindIII and NheI and ligated into the puromycin resistance cassette of pBabePuro-Ftractin-eGFP. Digested inserts and vectors were purified and ligated using T4 DNA Ligase (NEB). For bacterial transformation, Stbl3 cells were used. Plasmid isolation was performed with the PureLink HiPure Plasmid Filter Maxiprep Kit (Thermo Fisher Scientific). Insertion was confirmed by DNA sequencing.

Lentiviral pCDH-TRE-DUSP6 and pCDH-TRE-SPRY2 were generated as follows: Amplified *Dusp6* and *Spry2* fragments and empty pCDH-TRE-CRE vector were digested by PspXI and NotI and ligated using T4 DNA Ligase (NEB). For bacterial transformation, Stbl3 cells were used. Plasmid isolation was performed with the PureLink HiPure Plasmid Filter Maxiprep Kit (Thermo Fisher Scientific). Insertion was checked by DNA sequencing. The pCDH-TRE-CRE vector was generated by replacing the CMV promoter in pCDH-EF1-Cre (upstream of EF1-Cre, PMID 25936644) with the TRE3G promoter so that cloned cDNAs can be expressed in an rtTA/Doxycycline-dependent manner. The TRE3G promoter was amplified from the pTRE3G vector (Clontech) and inserted into SpeI/EcoRI sites of pCDH-EF1-Cre.

### Key resources table

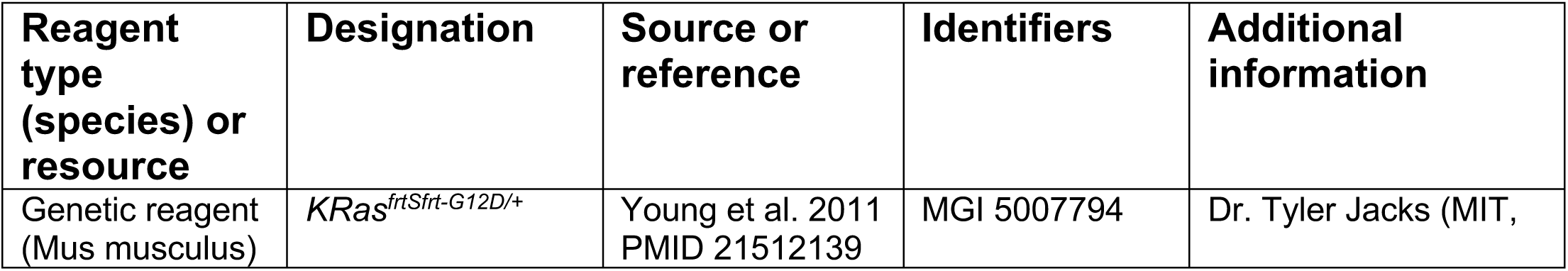
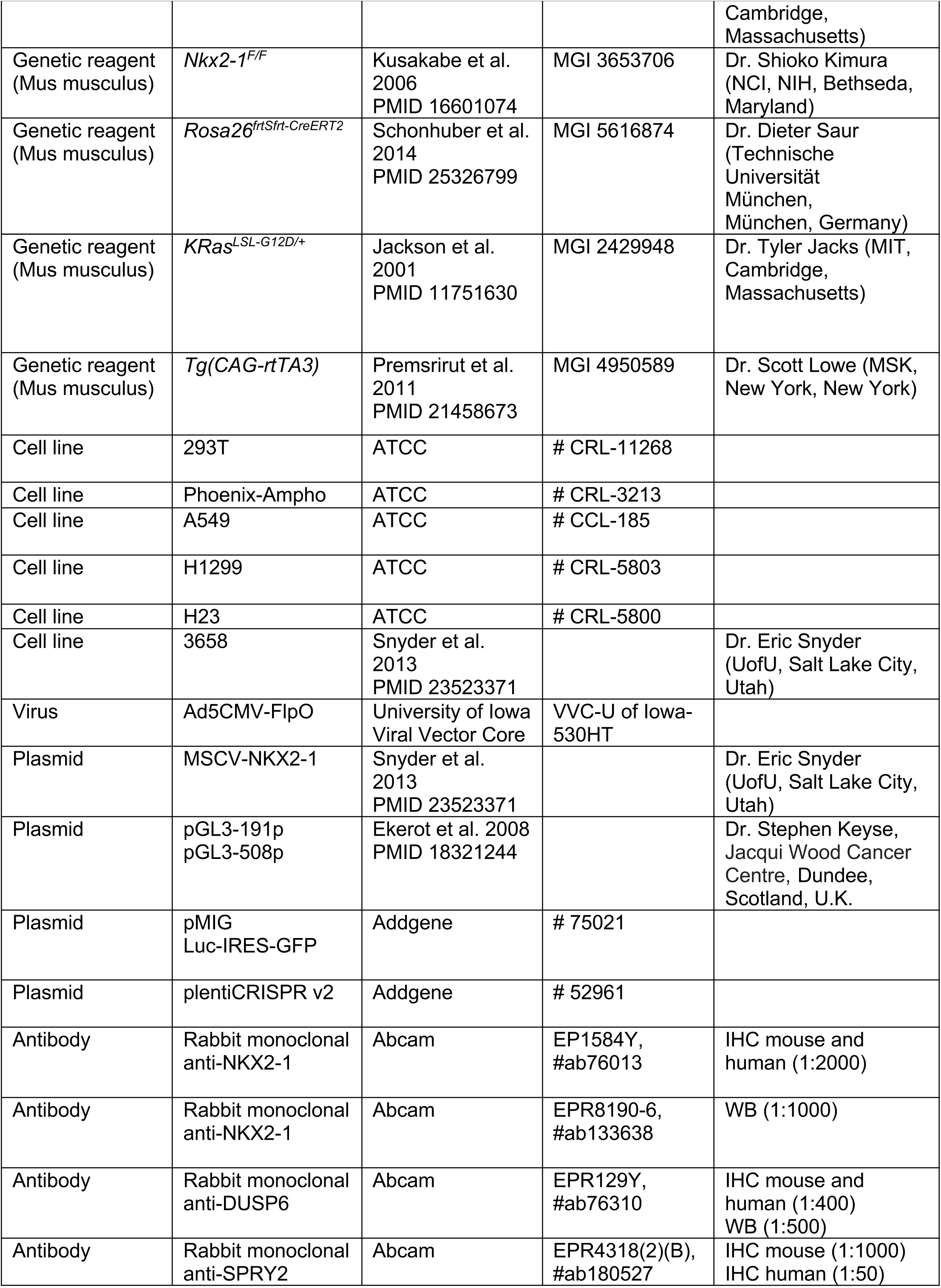
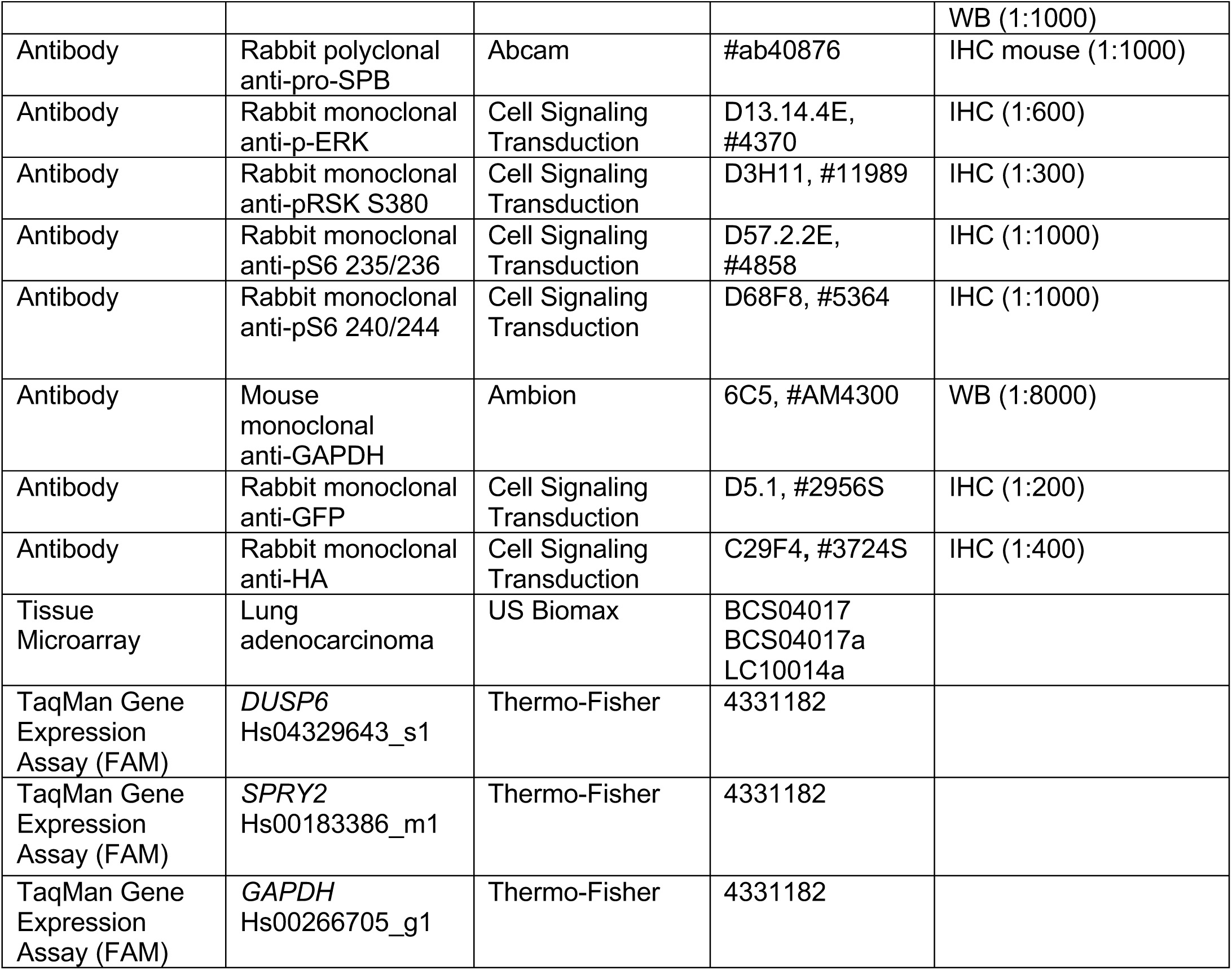

### Gene expression analysis

TCGA data for LUAD was not filtered for any demographic or clinical information. Upper quantile normalized FPKM values vs standard FPKM values were log transformed (natural log). Pearson correlation (r values) were generated using the base R function “cor”. Significance was determined using the base R function “cor.test”. Fitted lines were generated using the base R function “lm”.

### Cell lines and cell culture

A549, Phoenix-Ampho, H1299, H23, and 293T cells (ATCC) were cultured in DMEM/High Glucose medium (Gibco) with 5% FBS (Sigma). Cells were tested for mycoplasma every 3-4 months. Mycoplasma-negative cells were used for all experiments. 3658 cells were described previously (24, 50). Cell line identity was authenticated using STR analysis at the University of Utah DNA Sequencing Core. CRISPR/Cas9 knockout cells were generated by Cas9/CRISPR transfection, 2-day puromycin selection, dilution cloning, and sequencing.

### Retro- and lentivirus production and cell line infection

Retrovirus MSCV and MSCV-NKX2-1 and were produced by transfection of HEK293T cells with TransIT-293 (Mirus Bio, Madison, WI) with the retrovirus backbone as well as packaging vectors gag/pol and CMV-VSV-G. Transfections were carried-out in Opti-mem (Thermo Fisher Scientific) following Mirus’s TransIT-293 protocol. pBabe-H2B-mCherry-Ftractin-eGFP retrovirus was produced by transfection of Phoenix-Ampho cells. 24 hours after transfection, the media in the virus-producing cells was exchanged for DMEM with 30% FBS. Supernatant was collected at 48, 60, and 72 hr after transfection, filtered through a 0.45 μm filter, and used with 8 μg/ml polybrene to infect target cell lines once per day for three days. Following MSCV-NKX2-1 infection, cells were selected with 2.5 μg/ml puromycin for 1 week to achieve stable transduction. To generate H1299 cells expressing H2B-mCherry and NKX2-1, cells were first infected with pBabe-H2B-mcherry-Ftractin-EGFP retrovirus and sorted for mCherry and eGFP co-expression by flow cytometry. Positive cells were then infected with MSCV-EV or MSCV-NKX2.1 retrovirus and selected with puromycin. To generate the H1299 cells expressing luciferase, cells were infected with pMIG-luciferase-IRES-GFP and sorted for GFP expression. Cells infected with MSCV-EV or -NKX2-1 were maintained in 0.5 μg/ml puromycin prior to use for experiments.

Lentivirus pCW22-TRE-NKX2-1, pCDH-TRE-DUSP6, pCDH-TRE-SPRY2, and pMIG-luciferase-IRES-GFP was produced by transfection of 293T cells with TransIT-293 (Mirus Bio, Madison, WI), lentiviral backbone as well as packaging vectors Δ8.9 (gag/pol) packaging vector and CMV-VSV-G envelope vector. Supernatant was collected at 36, 48, 60 and 72 hrs after transfection. pCW22-TRE-NKX2-1 was used to infect A549 and H1299 DUSP6 and SPRY2 CRISPR knockout cells in the presence of 8 µg/ml polybrene for 24 hrs. Cells were selected with blasticidin for 1 week. NKX2-1 expression was induced in cells with 1 μg/ml of doxycycline and confirmed by Western blotting. Ultracentrifugation at 25,000 rpm for 2 hrs was necessary to concentrate pCDH-TRE-DUSP6 and pCDH-TRE-SPRY2 virus for *in vivo* infection.

### Fluorescence-activated cell sorting (FACS)

Cells were sorted for GFP/mCherry double positivity (H2B-mCherry-Ftractin-eGFP) or GFP positivity (Luciferase-IRES-GFP). Cells were suspended in DMEM, 5% FBS supplemented with penicillin/streptomycin (Pen/Strep) and sorted using a FacsAria cell sorter (BD Biosciences) with help from the UofU Flow Cytometry Core.

### Real-time PCR (qRT-PCR)

Total RNA was extracted using TRIzol with the PureLink RNA mini kit (Thermo Fisher Scientific). RNA was treated with on-column RNase-Free DNAse I digestion (Invitrogen). cDNA was synthesized from 500 ng of extracted RNA using the iScript reverse transcription Supermix (Bio-Rad). Real-time quantitative PCR was performed in 4 technical replicates with the SsoAdvanced Universal Probes Supermix (Bio-Rad) and detected with a Bio-Rad CFX Connect Real-Time PCR detection system. Gene expression was calculated relative to *GAPDH* (loading control) using the 2^-(dCt.x-average(dCt.control))^ method and normalized to the control group for graphing. TaqMan Assays used in this study are listed in table above.

### Immunoblot analysis (Western, WB)

Cells were washed once with cold PBS and then lysed on ice using RIPA buffer (10 mm Tris-Cl, pH 8.0, 1 mm EDTA, 0.5 mm EGTA, 1% Triton X-100, 0.1% sodium deoxycholate, 0.1% SDS) supplemented with 1 mM sodium orthovanadate, 1 mM phenylmethylsulfonyl fluoride, 5 μg/ml leupeptin, 5 μg/ml aprotinin, and 5 μg/ml pepstatin A. Protein extracts were clarified by centrifugation at 14,000 rpm for 10 min to remove cell debris and concentrations were measured with Pierce Coomassie Plus Protein Assay reagent (Thermo Fisher Scientific). Lysates were then subjected to SDS-PAGE gels (BioRad), and transferred to a Nitrocellulose membrane. Membranes were probed with primary antibodies against DUSP6 (EPR129Y, Abcam), SPRY2 (EPR4318(2)(B), Abcam), NKX2-1 (EPR8190-6, Abcam), and GAPDH and subsequently with DyLight fluorochrome-conjugated secondary antibodies (Thermo Fisher Scientific). Signal was visualized with the Odyssey CLx Imaging System (LI-COR). p-ERK Westerns were carried out in cells starved of serum for 24 hours and stimulated with epidermal growth factor (EGF, 50 ng/ml) or phorbal 12-myristate (PMA, 40 ng/ml).

### Luciferase assay

A549 control cells and cells re-expressing NKX2-1 were plated in a white 96-well polypropylene assay plate (Corning) at a density of 10,000 cells per well, 3 replicate wells per condition. Cells were transfected the next day with pRenilla (Addgene) and empty pGL3 vector or pGL3 vector containg the DUSP6 promoter regions 191p and 508p using Lipofectamine 2000 (Invitrogen). After 24 hours the Dual-Glo Luciferase Assay System (Promega) was used and luminescence was measured by the Synergy HT microplate reader (Biotek).

### Proliferation assay

Proliferation was assessed by either manual counting with trypan blue in triplicate or with Janus Green B (Sigma Aldrich #2869-83-2) with four technical replicates read on an Epoch2 (Biotek) plate reader at 620 nm.

### 2D migration assay and analysis

Cells were plated on acid-etched glass-bottomed 12-well plates (Matek). Two days after plating and just before imaging, medium was changed to Fluorobrite (Invitrogen) with 10% FBS and 20 mm HEPES and stained with DRAQ5 (ThermoFisher, 2 μM) to label the nuclei for tracking. Cell migration was imaged by phase contrast microscopy, at 37 °C, 5% CO_2_ on a Nikon Ti inverted microscope with a Plan Fluor ELWD 20× air objective and an environmental chamber, at the UofU Cell Imaging Core. Images were acquired with a Zyla cMOS camera using Elements every 10 mins over 15 h. Cells were tracked automatically in MATLAB using custom software based off of u-track multiple-particle tracking. The explored surface area explored by each tracked cell was calculated as the Mean Squared Displacement (MSD), the average square displacement between positions on a migration trajectory over increasing time intervals. Cell motion was tested for persistent random walk, in which the mean squared displacement increases in a superdiffusive manner: MSD(t) ∝ t^α^, where 1<α<2). Only cells exhibiting persistent random walk were included in the velocity and directionality calculations. Significance was calculated using the two-sample nonparametric Kolmogorov–Smirnov test at 5% significance, which compared the continuous, one-dimensional probability distributions.

### 3D invasion assay and analysis

H1299 cells were stably infected with pBabe-H2B-mCherry-Ftractin-eGFP and then infected with MSCV EV or NKX viral vectors. Spheroids were prepared with 3000 cells seeded in a nucleon Sphera U bottom plate (Thermo Scientific). After 3-4 days, spheroids were sandwiched between two 8 ul layers of a 2 mg/ml Collagen I gel in 15 well μ-slides (Ibidi, μ-slide angiogenesis #81501) (70). Reagents, plates, and tips were chilled to prevent early gelation. Collagen pre-gel was prepared with 4mg/ml Rat-tail collagen (Advanced Biomatrix) mixed 1:10 with 10x DMEM (Gibco), neutralized with 1 M NaOH to a pH of 7.5, and diluted to 2 mg/ml with 1x DMEM (Gibco). The first layer of Collagen was polymerized at 37°C for 10 minutes. Spheroids were suspended in 8 μl of collagen I pre-gel and incubated for 30 min at 37°C, media was added to cover the spheroid (1:1 DMEM:F12 phenol red free media (Gibco), with 10% FBS, 20 mM HEPES, and 1% Pen/Strep), and encapsulated spheroids were further incubated for 4-6 hours. Spheroids were imaged live on a Leica SP8 WLL confocal microscope equipped with live cell imaging chamber and Plan Apo PL APO 10X/0.40 objective (150 μm Z-stacks for a total Z of 2 μm). CO_2_ was maintained at 5% and temperature at 37°C. Images were acquired every 20 minutes for 14-18 hours. Cell invasion away from the spheroid was tracked live. Individual Cell tracks of cells that left the spheroid were analyzed using the Imaris “spots function” in Imaris, which identifies moving cells from background noise in motion (est. particle diameter 22 μm, quality > 1.5, autoregressive motion, max distance 20 μm, max Gap Size = 1, filtered for tracks greater than 1000 sec and track length greater than 50 μm to reduce noise). Erroneous tracks in the spheroid center were removed by including filters for Ar1Mean, Track Displacement Length, and Distance from image border. Any remaining broken tracks were corrected manually.

### Tumor xenograft and analysis

With the University of Utah PRR Core, IACUC #18-11004, subcutaneous tumors were generated and monitored as follows: 5 x10^6^ A549 cells, expressing MSCV-NKX2-1 were mixed with Matrigel and injected subcutaneously into the flank of NSG mice (both male and female mice). Tumors were resected and weighed after 6 weeks. Lungs were harvested, sectioned, stained by H&E (Core-ARUP look AP Histology Research), and examined manually by board-certified pathologist (E.L.S.) for the presence of micrometastases. For CRISPR knockouts, A549 cells with CRISPR-knockout of *Gfp* (control) or *Dusp6* were expressing both doxycycline (Dox)- inducible TRE-NKX2-1 and luciferase-IRES-GFP. Tumor burden was monitored weekly by caliper and bioluminescence measurements (IVIS). After reaching 150-400 mm^3^ tumor volumes, the mice were fed Dox (Envigo) for 4 weeks.

For the orthotopic tumors, H1299 cells were expressing luciferase-IRES-GFP and MSCV-NKX2-1, and 0.5×10^6^ cells were mixed with Matrigel and transplanted into the lungs of NOD-SCID mice by intrathoracic injection. Tumor burden was monitored weekly from bioluminescence by IVIS Spectrum and reported as background-corrected total flux, which is average radiance (flux per unit area and unit solid angle) integrated over the region of interest.

### Genetically engineered mice and tumor initiation

Under University of Utah IACUC #18-08005, mice harboring *KRas^frtSfrt-G12D/+^;Nkx2-1^f/f^; Rosa^frtSfrt-CreERT2^* alleles were infected intratracheally with 1 x 10^7^ pfu/mouse Ad5CMV-FlpO (University of Iowa Viral Vector Core). FlpO recombined Frt-STOP-Frt for simultaneous KRAS^G12D^ and Cre-ER expression. After 1 weeks, Tamoxifen was used to activate Cre^ERT2^ in established lesions and delete *Nkx2-1*. Tamoxifen was delivered by intraperitoneal injection of 120 mg/kg tamoxifen (Sigma), dissolved in corn oil. Mice received a total of 6 injections over the course of 9 days. Lungs were harvested 20 weeks after tumor initiation.

*Kras^LSL-G12D/+^; Nkx2-1^F/F^* mice and *Kras^LSL-G12D/+^; Nkx2-1^F/F^*; *CAG-rtTA3* mice were infected intratracheally with 4×10^4^ pfu/mouse lentivirus pCDH-TRE-DUSP6 or pCDH-TRE-SPRY2 that encoded 1) doxycycline inducible *Dusp6* or *Spry2* and 2) constitutive Cre. The *TRE* promoter requires the tetracycline-controlled transactivator for expression. In mice with the *CAG-rtTA3* transgene, ubiquitous expression of the third-generation *rtTA* reverse tetracycline-regulated transactivator gene repressed expression. The doxycycline diet (Envigo, 625 mg/kg dose) was administered for 1 week, which released the rtTA repression and induced exogenous DUSP6 or SPRY2 in established tumors. Tumor burden was quantified by histological analysis.

### Statistics

Significance’s calculated in MATLAB: two-sided Dunnett test, one way ANOVA with Tukey-Kramer post-hoc test, two-sample Kolmogorov-Smirnov (K-S), or Wilcox Mann-Whitney as appropriate. Correlation analyses and Pearson’s correlation coefficient calculated in R. Not significant (NS) for *p*>0.05, * *p*<0.05, ** *p*<0.01, *** *p*<0.001, **** p<0.0001.

## Acknowledgements

Thanks to Dr. Stephen Keyse for the gift of the DUSP6 promoter constructs and to Dr. Doug Mackay for the gift of H2B-mCherry-N2. Flow cytometry work was supported by the University of Utah Flow Cytometry Facility and the National Cancer Institute through 5P30CA042014-24 and by the National Center for Research Resources under 1S10RR026802-01. Cell imaging was supported by the University of Utah Cell Imaging Core. Xenografts were supported by the Huntsman Cancer Institute Preclinical Research Resource. M.C.M was supported by K01CA168850, an American Lung Association Research Grant, American Cancer Society RSG CSM130435, and R21CA215891.

## References

1. Pakkala S, Ramalingam SS. Personalized therapy for lung cancer: striking a moving target. JCI Insight. 2018;3(15). Epub 2018/08/10. doi: 10.1172/jci.insight.120858. PubMed PMID: 30089719; PMCID: PMC6129126.

2. Cancer Genome Atlas Research N. Comprehensive molecular profiling of lung adenocarcinoma. Nature. 2014;511(7511):543–50. doi: 10.1038/nature13385. PubMed PMID: 25079552.

3. Ding L, Getz G, Wheeler DA, Mardis ER, McLellan MD, Cibulskis K, Sougnez C, Greulich H, Muzny DM, Morgan MB, Fulton L, Fulton RS, Zhang Q, Wendl MC, Lawrence MS, Larson DE, Chen K, Dooling DJ, Sabo A, Hawes AC, Shen H, Jhangiani SN, Lewis LR, Hall O, Zhu Y, Mathew T, Ren Y, Yao J, Scherer SE, Clerc K, Metcalf GA, Ng B, Milosavljevic A, Gonzalez-Garay ML, Osborne JR, Meyer R, Shi X, Tang Y, Koboldt DC, Lin L, Abbott R, Miner TL, Pohl C, Fewell G, Haipek C, Schmidt H, Dunford-Shore BH, Kraja A, Crosby SD, Sawyer CS, Vickery T, Sander S, Robinson J, Winckler W, Baldwin J, Chirieac LR, Dutt A, Fennell T, Hanna M, Johnson BE, Onofrio RC, Thomas RK, Tonon G, Weir BA, Zhao X, Ziaugra L, Zody MC, Giordano T, Orringer MB, Roth JA, Spitz MR, Wistuba, II, Ozenberger B, Good PJ, Chang AC, Beer DG, Watson MA, Ladanyi M, Broderick S, Yoshizawa A, Travis WD, Pao W, Province MA, Weinstock GM, Varmus HE, Gabriel SB, Lander ES, Gibbs RA, Meyerson M, Wilson RK. Somatic mutations affect key pathways in lung adenocarcinoma. Nature. 2008;455(7216):1069–75. Epub 2008/10/25. doi: 10.1038/nature07423. PubMed PMID: 18948947; PMCID: PMC2694412.

4. Dankort D, Filenova E, Collado M, Serrano M, Jones K, McMahon M. A new mouse model to explore the initiation, progression, and therapy of BRAFV600E-induced lung tumors. Genes & development. 2007;21(4):379–84. Epub 2007/02/15. doi: 10.1101/gad.1516407. PubMed PMID: 17299132; PMCID: PMC1804325.

5. Johnson L, Mercer K, Greenbaum D, Bronson RT, Crowley D, Tuveson DA, Jacks T. Somatic activation of the K-ras oncogene causes early onset lung cancer in mice. Nature. 2001;410(6832):1111–6. doi: 10.1038/35074129. PubMed PMID: 11323676.

6. Blasco RB, Francoz S, Santamaria D, Canamero M, Dubus P, Charron J, Baccarini M, Barbacid M. c-Raf, but not B-Raf, is essential for development of K-Ras oncogene-driven non-small cell lung carcinoma. Cancer cell. 2011;19(5):652–63. Epub 2011/04/26. doi: 10.1016/j.ccr.2011.04.002. PubMed PMID: 21514245; PMCID: PMC4854330.

7. Trejo CL, Juan J, Vicent S, Sweet-Cordero A, McMahon M. MEK1/2 inhibition elicits regression of autochthonous lung tumors induced by KRASG12D or BRAFV600E. Cancer research. 2012;72(12):3048–59. doi: 10.1158/0008-5472.CAN-11-3649. PubMed PMID: 22511580; PMCID: 3393094.

8. Jackson EL, Olive KP, Tuveson DA, Bronson R, Crowley D, Brown M, Jacks T. The differential effects of mutant p53 alleles on advanced murine lung cancer. Cancer research. 2005;65(22):10280–8. doi: 10.1158/0008-5472.CAN-05-2193. PubMed PMID: 16288016.

9. Noguchi M. Stepwise progression of pulmonary adenocarcinoma--clinical and molecular implications. Cancer metastasis reviews. 2010;29(1):15–21. doi: 10.1007/s10555-010-9210-y. PubMed PMID: 20108111.

10. Vicent S, Lopez-Picazo JM, Toledo G, Lozano MD, Torre W, Garcia-Corchon C, Quero C, Soria JC, Martin-Algarra S, Manzano RG, Montuenga LM. ERK1/2 is activated in non-small-cell lung cancer and associated with advanced tumours. Br J Cancer. 2004;90(5):1047–52. Epub 2004/03/05. doi: 10.1038/sj.bjc.6601644. PubMed PMID: 14997206; PMCID: PMC2409626.

11. Feldser DM, Kostova KK, Winslow MM, Taylor SE, Cashman C, Whittaker CA, Sanchez-Rivera FJ, Resnick R, Bronson R, Hemann MT, Jacks T. Stage-specific sensitivity to p53 restoration during lung cancer progression. Nature. 2010;468(7323):572–5. doi: 10.1038/nature09535. PubMed PMID: 21107428; PMCID: 3003305.

12. Junttila MR, Karnezis AN, Garcia D, Madriles F, Kortlever RM, Rostker F, Brown Swigart L, Pham DM, Seo Y, Evan GI, Martins CP. Selective activation of p53-mediated tumour suppression in high-grade tumours. Nature. 2010;468(7323):567–71. doi: 10.1038/nature09526. PubMed PMID: 21107427; PMCID: PMC3011233.

13. Zheng S, El-Naggar AK, Kim ES, Kurie JM, Lozano G. A genetic mouse model for metastatic lung cancer with gender differences in survival. Oncogene. 2007;26(48):6896–904. doi: 10.1038/sj.onc.1210493. PubMed PMID: 17486075.

14. Gilbert-Ross M, Konen J, Koo J, Shupe J, Robinson BS, Wiles WGt, Huang C, Martin WD, Behera M, Smith GH, Hill CE, Rossi MR, Sica GL, Rupji M, Chen Z, Kowalski J, Kasinski AL, Ramalingam SS, Fu H, Khuri FR, Zhou W, Marcus AI. Targeting adhesion signaling in KRAS, LKB1 mutant lung adenocarcinoma. JCI Insight. 2017;2(5):e90487. Epub 2017/03/16. doi: 10.1172/jci.insight.90487. PubMed PMID: 28289710; PMCID: PMC5333956 exists.

15. Heidorn SJ, Milagre C, Whittaker S, Nourry A, Niculescu-Duvas I, Dhomen N, Hussain J, Reis-Filho JS, Springer CJ, Pritchard C, Marais R. Kinase-dead BRAF and oncogenic RAS cooperate to drive tumor progression through CRAF. Cell. 2010;140(2):209–21. Epub 2010/02/10. doi: 10.1016/j.cell.2009.12.040. PubMed PMID: 20141835; PMCID: PMC2872605.

16. Nieto P, Ambrogio C, Esteban-Burgos L, Gomez-Lopez G, Blasco MT, Yao Z, Marais R, Rosen N, Chiarle R, Pisano DG, Barbacid M, Santamaria D. A Braf kinase-inactive mutant induces lung adenocarcinoma. Nature. 2017;548(7666):239–43. Epub 2017/08/08. doi: 10.1038/nature23297. PubMed PMID: 28783725; PMCID: PMC5648056.

17. Cicchini M, Buza EL, Sagal KM, Gudiel AA, Durham AC, Feldser DM. Context-Dependent Effects of Amplified MAPK Signaling during Lung Adenocarcinoma Initiation and Progression. Cell reports. 2017;18(8):1958–69. Epub 2017/02/24. doi: 10.1016/j.celrep.2017.01.069. PubMed PMID: 28228261; PMCID: PMC5405440.

18. Shaw AT, Meissner A, Dowdle JA, Crowley D, Magendantz M, Ouyang C, Parisi T, Rajagopal J, Blank LJ, Bronson RT, Stone JR, Tuveson DA, Jaenisch R, Jacks T. Sprouty-2 regulates oncogenic K-ras in lung development and tumorigenesis. Genes & development. 2007;21(6):694–707. doi: 10.1101/gad.1526207. PubMed PMID: 17369402; PMCID: 1820943.

19. Wagner PL, Perner S, Rickman DS, LaFargue CJ, Kitabayashi N, Johnstone SF, Weir BA, Meyerson M, Altorki NK, Rubin MA. In situ evidence of KRAS amplification and association with increased p21 levels in non-small cell lung carcinoma. Am J Clin Pathol. 2009;132(4):500–5. Epub 2009/09/19. doi: 10.1309/AJCPF10ZUNSOLIFG. PubMed PMID: 19762526.

20. Zhang Z, Kobayashi S, Borczuk AC, Leidner RS, Laframboise T, Levine AD, Halmos B. Dual specificity phosphatase 6 (DUSP6) is an ETS-regulated negative feedback mediator of oncogenic ERK signaling in lung cancer cells. Carcinogenesis. 2010;31(4):577–86. doi: 10.1093/carcin/bgq020. PubMed PMID: 20097731; PMCID: 2847094.

21. Travis WD, Brambilla E, Noguchi M, Nicholson AG, Geisinger K, Yatabe Y, Powell CA, Beer D, Riely G, Garg K, Austin JH, Rusch VW, Hirsch FR, Jett J, Yang PC, Gould M. International Association for the Study of Lung Cancer/American Thoracic Society/European Respiratory Society: international multidisciplinary classification of lung adenocarcinoma: executive summary. Proc Am Thorac Soc. 2011;8(5):381–5. Epub 2011/09/20. doi: 10.1513/pats.201107-042ST. PubMed PMID: 21926387.

22. Russell PA, Wainer Z, Wright GM, Daniels M, Conron M, Williams RA. Does lung adenocarcinoma subtype predict patient survival?: A clinicopathologic study based on the new International Association for the Study of Lung Cancer/American Thoracic Society/European Respiratory Society international multidisciplinary lung adenocarcinoma classification. J Thorac Oncol. 2011;6(9):1496–504. Epub 2011/06/07. doi: 10.1097/JTO.0b013e318221f701. PubMed PMID: 21642859.

23. Yoshizawa A, Motoi N, Riely GJ, Sima CS, Gerald WL, Kris MG, Park BJ, Rusch VW, Travis WD. Impact of proposed IASLC/ATS/ERS classification of lung adenocarcinoma: prognostic subgroups and implications for further revision of staging based on analysis of 514 stage I cases. Mod Pathol. 2011;24(5):653–64. Epub 2011/01/22. doi: 10.1038/modpathol.2010.232. PubMed PMID: 21252858.

24. Snyder EL, Watanabe H, Magendantz M, Hoersch S, Chen TA, Wang DG, Crowley D, Whittaker CA, Meyerson M, Kimura S, Jacks T. Nkx2-1 represses a latent gastric differentiation program in lung adenocarcinoma. Molecular cell. 2013;50(2):185–99. doi: 10.1016/j.molcel.2013.02.018. PubMed PMID: 23523371; PMCID: 3721642.

25. Winslow MM, Dayton TL, Verhaak RG, Kim-Kiselak C, Snyder EL, Feldser DM, Hubbard DD, DuPage MJ, Whittaker CA, Hoersch S, Yoon S, Crowley D, Bronson RT, Chiang DY, Meyerson M, Jacks T. Suppression of lung adenocarcinoma progression by Nkx2-1. Nature. 2011;473(7345):101–4. doi: 10.1038/nature09881. PubMed PMID: 21471965; PMCID: 3088778.

26. Marjanovic ND, Hofree M, Chan JE, Canner D, Wu K, Trakala M, Hartmann GG, Smith OC, Kim JY, Evans KV, Hudson A, Ashenberg O, Porter CBM, Bejnood A, Subramanian A, Pitter K, Yan Y, Delorey T, Phillips DR, Shah N, Chaudhary O, Tsankov A, Hollmann T, Rekhtman N, Massion PP, Poirier JT, Mazutis L, Li R, Lee JH, Amon A, Rudin CM, Jacks T, Regev A, Tammela T. Emergence of a High-Plasticity Cell State during Lung Cancer Evolution. Cancer cell. 2020;38(2):229–46 e13. Epub 2020/07/25. doi: 10.1016/j.ccell.2020.06.012. PubMed PMID: 32707077; PMCID: PMC7745838.

27. Maeda Y, Tsuchiya T, Hao H, Tompkins DH, Xu Y, Mucenski ML, Du L, Keiser AR, Fukazawa T, Naomoto Y, Nagayasu T, Whitsett JA. Kras(G12D) and Nkx2-1 haploinsufficiency induce mucinous adenocarcinoma of the lung. The Journal of clinical investigation. 2012;122(12):4388–400. doi: 10.1172/JCI64048. PubMed PMID: 23143308; PMCID: 3533546.

28. Barletta JA, Perner S, Iafrate AJ, Yeap BY, Weir BA, Johnson LA, Johnson BE, Meyerson M, Rubin MA, Travis WD, Loda M, Chirieac LR. Clinical significance of TTF-1 protein expression and TTF-1 gene amplification in lung adenocarcinoma. Journal of cellular and molecular medicine. 2009;13(8B):1977–86. doi: 10.1111/j.1582-4934.2008.00594.x. PubMed PMID: 19040416; PMCID: 2830395.

29. Berghmans T, Paesmans M, Mascaux C, Martin B, Meert AP, Haller A, Lafitte JJ, Sculier JP. Thyroid transcription factor 1--a new prognostic factor in lung cancer: a meta-analysis. Annals of oncology : official journal of the European Society for Medical Oncology / ESMO. 2006;17(11):1673–6. doi: 10.1093/annonc/mdl287. PubMed PMID: 16980598.

30. Cardnell RJ, Behrens C, Diao L, Fan Y, Tang X, Tong P, Minna JD, Mills GB, Heymach JV, Wistuba, II, Wang J, Byers LA. An Integrated Molecular Analysis of Lung Adenocarcinomas Identifies Potential Therapeutic Targets among TTF1-Negative Tumors, Including DNA Repair Proteins and Nrf2. Clinical cancer research : an official journal of the American Association for Cancer Research. 2015;21(15):3480–91. Epub 2015/04/17. doi: 10.1158/1078-0432.CCR-14-3286. PubMed PMID: 25878335; PMCID: PMC4526428.

31. Gillies TE, Pargett M, Silva JM, Teragawa C, McCormick F, Albeck JG. Oncogenic mutant RAS signaling activity is rescaled by the ERK/MAPK pathway. bioRxiv. 2020:2020.02.17.952093. doi: 10.1101/2020.02.17.952093.

32. Avraham R, Yarden Y. Feedback regulation of EGFR signalling: decision making by early and delayed loops. Nature reviews Molecular cell biology. 2011;12(2):104–17. Epub 2011/01/22. doi: 10.1038/nrm3048. PubMed PMID: 21252999.

33. Lake D, Correa SA, Muller J. Negative feedback regulation of the ERK1/2 MAPK pathway. Cell Mol Life Sci. 2016;73(23):4397–413. Epub 2016/10/23. doi: 10.1007/s00018-016-2297-8. PubMed PMID: 27342992; PMCID: PMC5075022.

34. Stern DF. Keeping Tumors Out of the MAPK Fitness Zone. Cancer discovery. 2018;8(1):20–3. Epub 2018/01/10. doi: 10.1158/2159-8290.CD-17-1243. PubMed PMID: 29311225.

35. Nunes-Xavier C, Roma-Mateo C, Rios P, Tarrega C, Cejudo-Marin R, Tabernero L, Pulido R. Dual-specificity MAP kinase phosphatases as targets of cancer treatment. Anti-cancer agents in medicinal chemistry. 2011;11(1):109–32. PubMed PMID: 21288197.

36. Kidger AM, Keyse SM. The regulation of oncogenic Ras/ERK signalling by dual-specificity mitogen activated protein kinase phosphatases (MKPs). Semin Cell Dev Biol. 2016;50:125–32. Epub 2016/01/23. doi: 10.1016/j.semcdb.2016.01.009. PubMed PMID: 26791049; PMCID: PMC5056954.

37. Muda M, Theodosiou A, Rodrigues N, Boschert U, Camps M, Gillieron C, Davies K, Ashworth A, Arkinstall S. The dual specificity phosphatases M3/6 and MKP-3 are highly selective for inactivation of distinct mitogen-activated protein kinases. The Journal of biological chemistry. 1996;271(44):27205–8. Epub 1996/11/01. doi: 10.1074/jbc.271.44.27205. PubMed PMID: 8910287.

38. Unni AM, Harbourne B, Oh MH, Wild S, Ferrarone JR, Lockwood WW, Varmus H. Hyperactivation of ERK by multiple mechanisms is toxic to RTK-RAS mutation-driven lung adenocarcinoma cells. Elife. 2018;7. Epub 2018/11/27. doi: 10.7554/eLife.33718. PubMed PMID: 30475204; PMCID: PMC6298772.

39. Chen HY, Yu SL, Chen CH, Chang GC, Chen CY, Yuan A, Cheng CL, Wang CH, Terng HJ, Kao SF, Chan WK, Li HN, Liu CC, Singh S, Chen WJ, Chen JJ, Yang PC. A five-gene signature and clinical outcome in non-small-cell lung cancer. N Engl J Med. 2007;356(1):11–20. Epub 2007/01/05. doi: 10.1056/NEJMoa060096. PubMed PMID: 17202451.

40. Okudela K, Yazawa T, Woo T, Sakaeda M, Ishii J, Mitsui H, Shimoyamada H, Sato H, Tajiri M, Ogawa N, Masuda M, Takahashi T, Sugimura H, Kitamura H. Down-regulation of DUSP6 expression in lung cancer: its mechanism and potential role in carcinogenesis. Am J Pathol. 2009;175(2):867–81. Epub 2009/07/18. doi: 10.2353/ajpath.2009.080489. PubMed PMID: 19608870; PMCID: PMC2716981.

41. Hrustanovic G, Olivas V, Pazarentzos E, Tulpule A, Asthana S, Blakely CM, Okimoto RA, Lin L, Neel DS, Sabnis A, Flanagan J, Chan E, Varella-Garcia M, Aisner DL, Vaishnavi A, Ou SH, Collisson EA, Ichihara E, Mack PC, Lovly CM, Karachaliou N, Rosell R, Riess JW, Doebele RC, Bivona TG. RAS-MAPK dependence underlies a rational polytherapy strategy in EML4-ALK-positive lung cancer. Nature medicine. 2015;21(9):1038–47. Epub 2015/08/25. doi: 10.1038/nm.3930. PubMed PMID: 26301689; PMCID: PMC4734742.

42. Mason JM, Morrison DJ, Basson MA, Licht JD. Sprouty proteins: multifaceted negative-feedback regulators of receptor tyrosine kinase signaling. Trends in cell biology. 2006;16(1):45–54. doi: 10.1016/j.tcb.2005.11.004. PubMed PMID: 16337795.

43. Sutterluty H, Mayer CE, Setinek U, Attems J, Ovtcharov S, Mikula M, Mikulits W, Micksche M, Berger W. Down-regulation of Sprouty2 in non-small cell lung cancer contributes to tumor malignancy via extracellular signal-regulated kinase pathway-dependent and -independent mechanisms. Mol Cancer Res. 2007;5(5):509–20. Epub 2007/05/19. doi: 10.1158/1541-7786.MCR-06-0273. PubMed PMID: 17510316.

44. Takeuchi T, Tomida S, Yatabe Y, Kosaka T, Osada H, Yanagisawa K, Mitsudomi T, Takahashi T. Expression profile-defined classification of lung adenocarcinoma shows close relationship with underlying major genetic changes and clinicopathologic behaviors. Journal of clinical oncology : official journal of the American Society of Clinical Oncology. 2006;24(11):1679–88. Epub 2006/03/22. doi: 10.1200/JCO.2005.03.8224. PubMed PMID: 16549822.

45. Yatabe Y, Kosaka T, Takahashi T, Mitsudomi T. EGFR mutation is specific for terminal respiratory unit type adenocarcinoma. Am J Surg Pathol. 2005;29(5):633–9. Epub 2005/04/16. PubMed PMID: 15832087.

46. Watanabe H, Francis JM, Woo MS, Etemad B, Lin W, Fries DF, Peng S, Snyder EL, Tata PR, Izzo F, Schinzel AC, Cho J, Hammerman PS, Verhaak RG, Hahn WC, Rajagopal J, Jacks T, Meyerson M. Integrated cistromic and expression analysis of amplified NKX2-1 in lung adenocarcinoma identifies LMO3 as a functional transcriptional target. Genes & development. 2013;27(2):197–210. Epub 2013/01/17. doi: 10.1101/gad.203208.112. PubMed PMID: 23322301; PMCID: 3566312.

47. Ding W, Bellusci S, Shi W, Warburton D. Functional analysis of the human Sprouty2 gene promoter. Gene. 2003;322:175–85. Epub 2003/12/04. doi: 10.1016/j.gene.2003.09.004. PubMed PMID: 14644509.

48. Ekerot M, Stavridis MP, Delavaine L, Mitchell MP, Staples C, Owens DM, Keenan ID, Dickinson RJ, Storey KG, Keyse SM. Negative-feedback regulation of FGF signalling by DUSP6/MKP-3 is driven by ERK1/2 and mediated by Ets factor binding to a conserved site within the DUSP6/MKP-3 gene promoter. Biochem J. 2008;412(2):287–98. Epub 2008/03/07. doi: 10.1042/BJ20071512. PubMed PMID: 18321244; PMCID: PMC2474557.

49. Gross I, Armant O, Benosman S, de Aguilar JL, Freund JN, Kedinger M, Licht JD, Gaiddon C, Loeffler JP. Sprouty2 inhibits BDNF-induced signaling and modulates neuronal differentiation and survival. Cell Death Differ. 2007;14(10):1802–12. Epub 2007/06/30. doi: 10.1038/sj.cdd.4402188. PubMed PMID: 17599098.

50. Samson SC, Elliott A, Mueller BD, Kim Y, Carney KR, Bergman JP, Blenis J, Mendoza MC. p90 ribosomal S6 kinase (RSK) phosphorylates myosin phosphatase and thereby controls edge dynamics during cell migration. The Journal of biological chemistry. 2019;294(28):10846–62. Epub 2019/05/30. doi: 10.1074/jbc.RA119.007431. PubMed PMID: 31138649; PMCID: PMC6635457.

51. Premsrirut PK, Dow LE, Kim SY, Camiolo M, Malone CD, Miething C, Scuoppo C, Zuber J, Dickins RA, Kogan SC, Shroyer KR, Sordella R, Hannon GJ, Lowe SW. A rapid and scalable system for studying gene function in mice using conditional RNA interference. Cell. 2011;145(1):145–58. Epub 2011/04/05. doi: 10.1016/j.cell.2011.03.012. PubMed PMID: 21458673; PMCID: PMC3244080.

52. Burotto M, Chiou VL, Lee JM, Kohn EC. The MAPK pathway across different malignancies: a new perspective. Cancer. 2014;120(22):3446–56. doi: 10.1002/cncr.28864. PubMed PMID: 24948110; PMCID: PMC4221543.

53. Vartanian S, Bentley C, Brauer MJ, Li L, Shirasawa S, Sasazuki T, Kim JS, Haverty P, Stawiski E, Modrusan Z, Waldman T, Stokoe D. Identification of mutant K-Ras-dependent phenotypes using a panel of isogenic cell lines. The Journal of biological chemistry. 2013;288(4):2403–13. Epub 2012/11/29. doi: 10.1074/jbc.M112.394130. PubMed PMID: 23188824; PMCID: PMC3554910.

54. Consortium APG. AACR Project GENIE: Powering Precision Medicine through an International Consortium. Cancer discovery. 2017;7(8):818–31. Epub 2017/06/03. doi: 10.1158/2159-8290.CD-17-0151. PubMed PMID: 28572459; PMCID: PMC5611790.

55. Kimura S, Hara Y, Pineau T, Fernandez-Salguero P, Fox CH, Ward JM, Gonzalez FJ. The T/ebp null mouse: thyroid-specific enhancer-binding protein is essential for the organogenesis of the thyroid, lung, ventral forebrain, and pituitary. Genes & development. 1996;10(1):60–9. Epub 1996/01/01. doi: 10.1101/gad.10.1.60. PubMed PMID: 8557195.

56. Sutherland KD, Song JY, Kwon MC, Proost N, Zevenhoven J, Berns A. Multiple cells-of-origin of mutant K-Ras-induced mouse lung adenocarcinoma. Proceedings of the National Academy of Sciences of the United States of America. 2014;111(13):4952–7. Epub 2014/03/04. doi: 10.1073/pnas.1319963111. PubMed PMID: 24586047; PMCID: PMC3977239.

57. Xu X, Rock JR, Lu Y, Futtner C, Schwab B, Guinney J, Hogan BL, Onaitis MW. Evidence for type II cells as cells of origin of K-Ras-induced distal lung adenocarcinoma. Proceedings of the National Academy of Sciences of the United States of America. 2012;109(13):4910–5. Epub 2012/03/14. doi: 10.1073/pnas.1112499109. PubMed PMID: 22411819; PMCID: PMC3323959.

58. Diaz-Garcia CV, Agudo-Lopez A, Perez C, Prieto-Garcia E, Iglesias L, Ponce S, Rodriguez Garzotto A, Rodriguez-Peralto JL, Cortes-Funes H, Lopez-Martin JA, Agullo-Ortuno MT. Prognostic value of dual-specificity phosphatase 6 expression in non-small cell lung cancer. Tumour Biol. 2015;36(2):1199–206. doi: 10.1007/s13277-014-2729-8. PubMed PMID: 25344212.

59. Piya S, Kim JY, Bae J, Seol DW, Moon AR, Kim TH. DUSP6 is a novel transcriptional target of p53 and regulates p53-mediated apoptosis by modulating expression levels of Bcl-2 family proteins. FEBS letters. 2012;586(23):4233–40. doi: 10.1016/j.febslet.2012.10.031. PubMed PMID: 23108049.

60. Ahmad MK, Abdollah NA, Shafie NH, Yusof NM, Razak SRA. Dual-specificity phosphatase 6 (DUSP6): a review of its molecular characteristics and clinical relevance in cancer. Cancer Biol Med. 2018;15(1):14–28. Epub 2018/03/17. doi: 10.20892/j.issn.2095-3941.2017.0107. PubMed PMID: 29545965; PMCID: PMC5842331.

61. Chiba I, Takahashi T, Nau MM, D’Amico D, Curiel DT, Mitsudomi T, Buchhagen DL, Carbone D, Piantadosi S, Koga H, et al. Mutations in the p53 gene are frequent in primary, resected non-small cell lung cancer. Lung Cancer Study Group. Oncogene. 1990;5(10):1603–10. Epub 1990/10/01. PubMed PMID: 1979160.

62. Ramkissoon A, Chaney KE, Milewski D, Williams KB, Williams RL, Choi K, Miller A, Kalin TV, Pressey JG, Szabo S, Azam M, Largaespada DA, Ratner N. Targeted Inhibition of the Dual Specificity Phosphatases DUSP1 and DUSP6 Suppress MPNST Growth via JNK. Clinical cancer research : an official journal of the American Association for Cancer Research. 2019;25(13):4117–27. Epub 2019/04/03. doi: 10.1158/1078-0432.CCR-18-3224. PubMed PMID: 30936125; PMCID: PMC6606396.

63. Shojaee S, Caeser R, Buchner M, Park E, Swaminathan S, Hurtz C, Geng H, Chan LN, Klemm L, Hofmann WK, Qiu YH, Zhang N, Coombes KR, Paietta E, Molkentin J, Koeffler HP, Willman CL, Hunger SP, Melnick A, Kornblau SM, Muschen M. Erk Negative Feedback Control Enables Pre-B Cell Transformation and Represents a Therapeutic Target in Acute Lymphoblastic Leukemia. Cancer cell. 2015;28(1):114–28. Epub 2015/06/16. doi: 10.1016/j.ccell.2015.05.008. PubMed PMID: 26073130; PMCID: PMC4565502.

64. Zewdu R, Mehrabad EM, Ingram K, Jones A, Camolotto SA, Mendoza MC, Spike B, Snyder EL. An NKX2-1/ERK/WNT feedback loop modulates gastric identity and response to targeted therapy in lung adenocarcinoma. bioRxiv. 2020:2020.02.25.965004. doi: 10.1101/2020.02.25.965004.

65. Kong XJ, Kuilman T, Shahrabi A, Oshuizen JB, Kemper K, Song JY, Niessen HWM, Rozeman EA, Foppen MHG, Lank CUB, Peeper DS. Cancer drug addiction is relayed by an ERK2-dependent phenotype switch. Nature. 2017;550(7675):270-+. doi: 10.1038/nature24037. PubMed PMID: WOS:000412829500051.

66. Davies AE, Pargett M, Siebert S, Gillies TE, Choi Y, Tobin SJ, Ram AR, Murthy V, Juliano C, Quon G, Bissell MJ, Albeck JG. Systems-Level Properties of EGFR-RAS-ERK Signaling Amplify Local Signals to Generate Dynamic Gene Expression Heterogeneity. Cell Syst. 2020;11(2):161–75 e5. Epub 2020/07/30. doi: 10.1016/j.cels.2020.07.004. PubMed PMID: 32726596; PMCID: PMC7856305.

67. Wu QN, Liao YF, Lu YX, Wang Y, Lu JH, Zeng ZL, Huang QT, Sheng H, Yun JP, Xie D, Ju HQ, Xu RH. Pharmacological inhibition of DUSP6 suppresses gastric cancer growth and metastasis and overcomes cisplatin resistance. Cancer Lett. 2018;412:243–55. Epub 2017/10/21. doi: 10.1016/j.canlet.2017.10.007. PubMed PMID: 29050982.

68. James NE, Beffa L, Oliver MT, Borgstadt AD, Emerson JB, Chichester CO, Yano N, Freiman RN, DiSilvestro PA, Ribeiro JR. Inhibition of DUSP6 sensitizes ovarian cancer cells to chemotherapeutic agents via regulation of ERK signaling response genes. Oncotarget. 2019;10(36):3315–27. Epub 2019/06/06. PubMed PMID: 31164954; PMCID: PMC6534361.

69. Zhang Y, Liu T, Meyer CA, Eeckhoute J, Johnson DS, Bernstein BE, Nusbaum C, Myers RM, Brown M, Li W, Liu XS. Model-based analysis of ChIP-Seq (MACS). Genome biology. 2008;9(9):R137. Epub 2008/09/19. doi: 10.1186/gb-2008-9-9-r137. PubMed PMID: 18798982; PMCID: PMC2592715.

70. Konen J, Summerbell E, Dwivedi B, Galior K, Hou Y, Rusnak L, Chen A, Saltz J, Zhou W, Boise LH, Vertino P, Cooper L, Salaita K, Kowalski J, Marcus AI. Image-guided genomics of phenotypically heterogeneous populations reveals vascular signalling during symbiotic collective cancer invasion. Nat Commun. 2017;8:15078. Epub 2017/05/13. doi: 10.1038/ncomms15078. PubMed PMID: 28497793; PMCID: PMC5437311.

