## Supplementary figures and images for "NKX2-1 controls lung cancer progression by inducing DUSP6 to dampen ERK activity"

### Supplemental Figures S1-S5

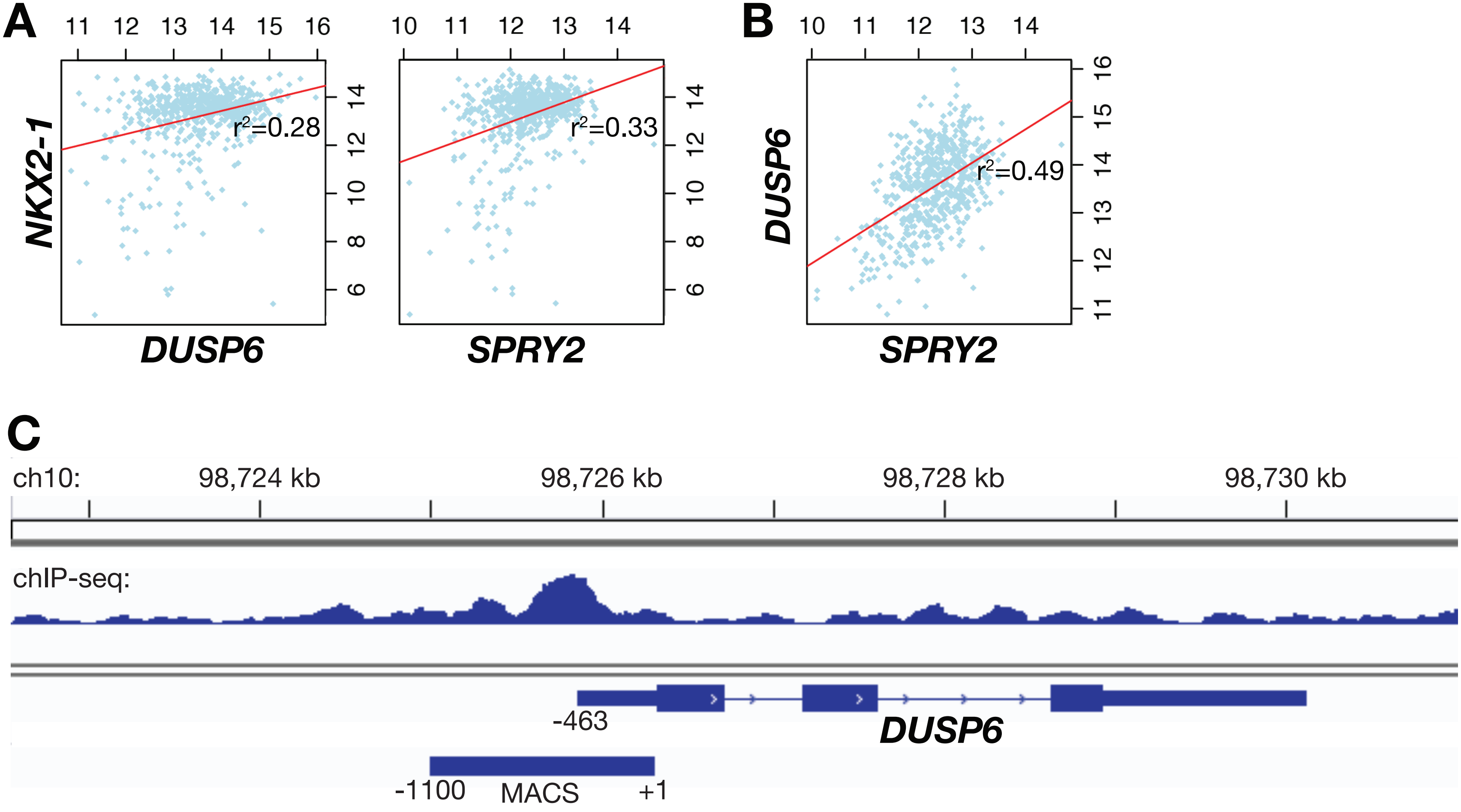

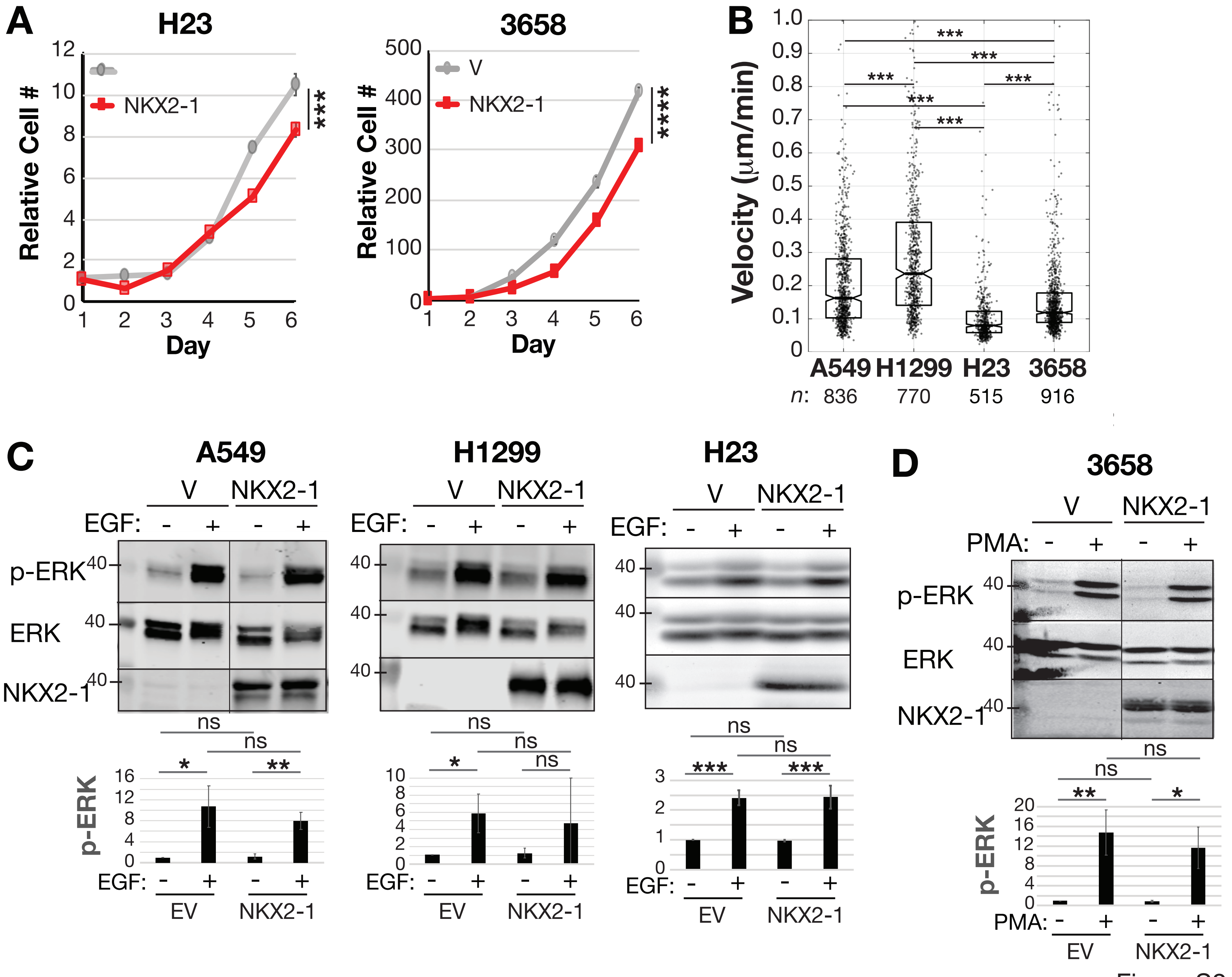

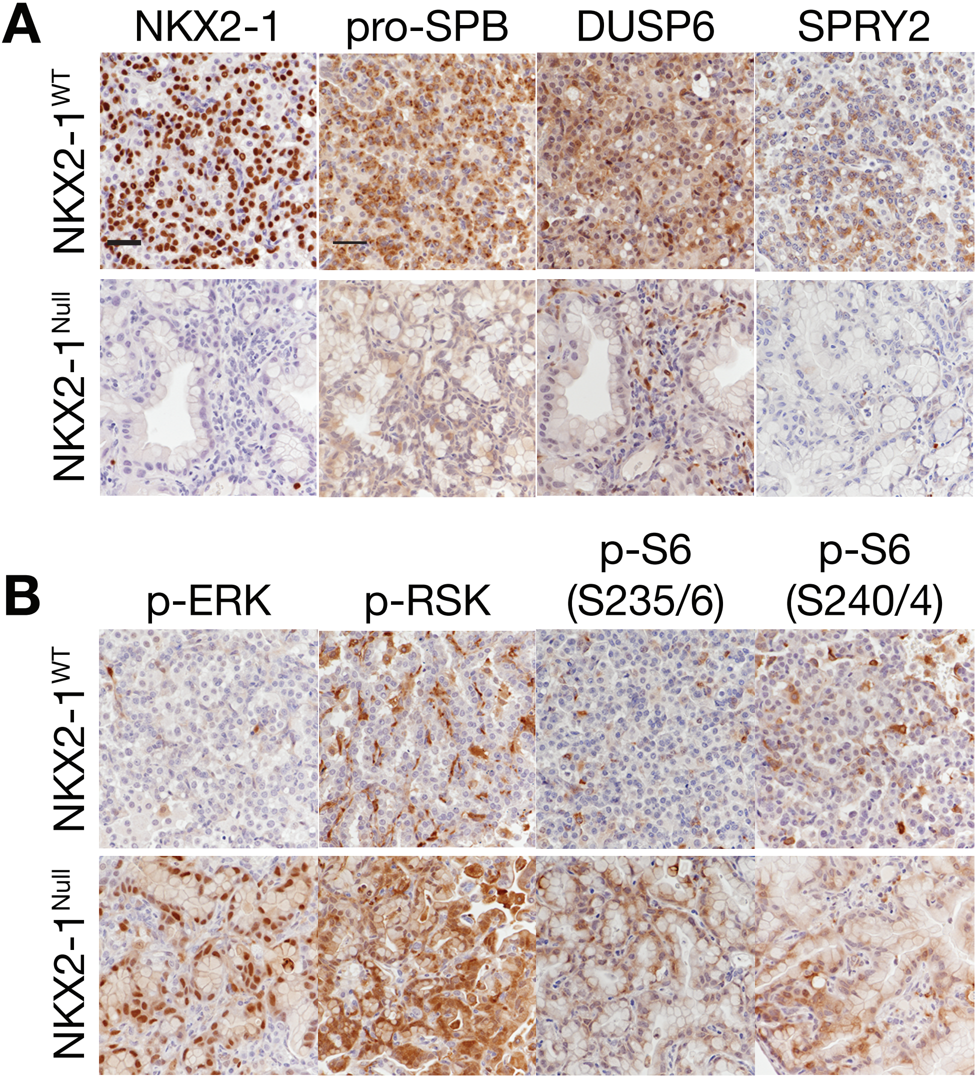

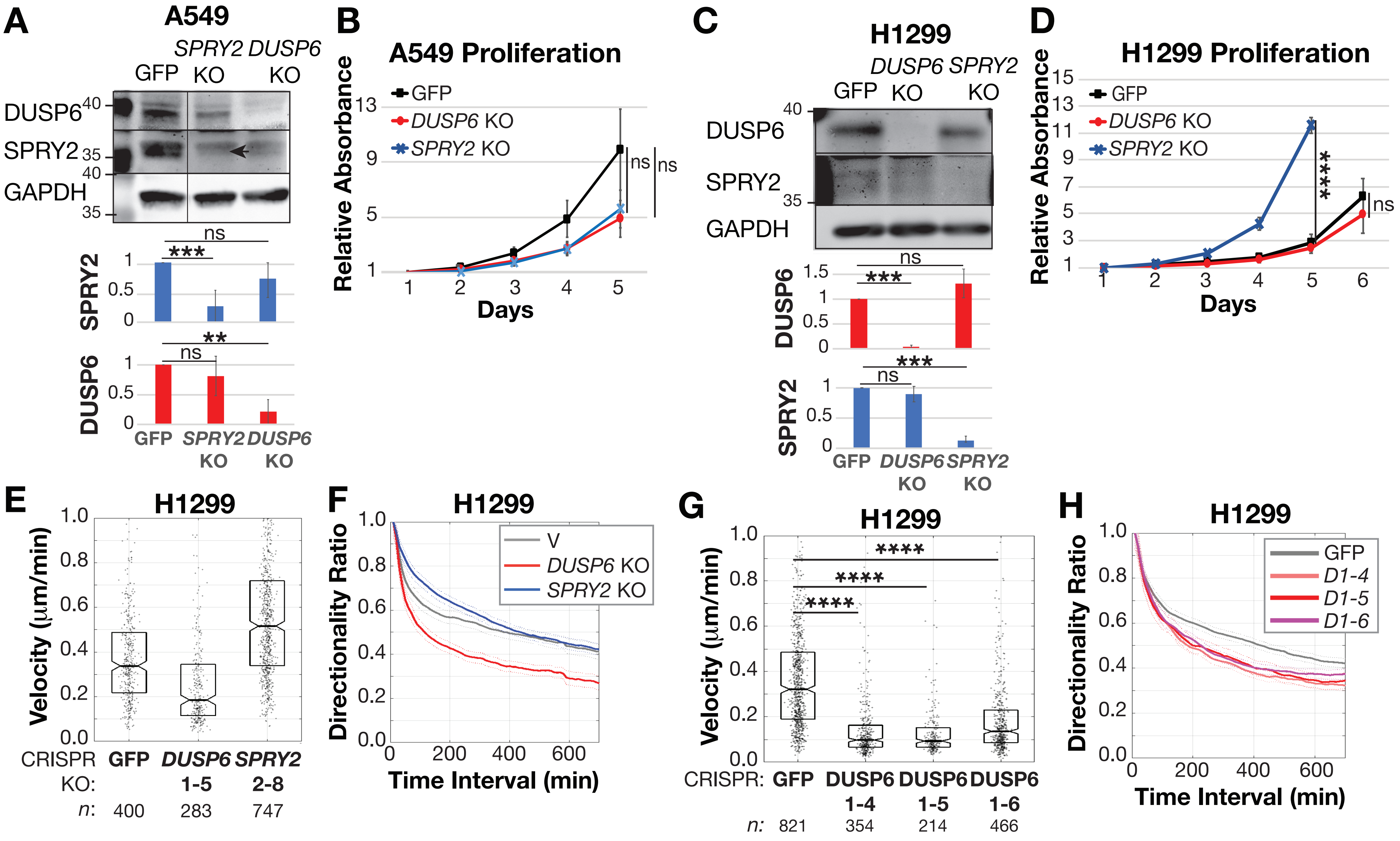

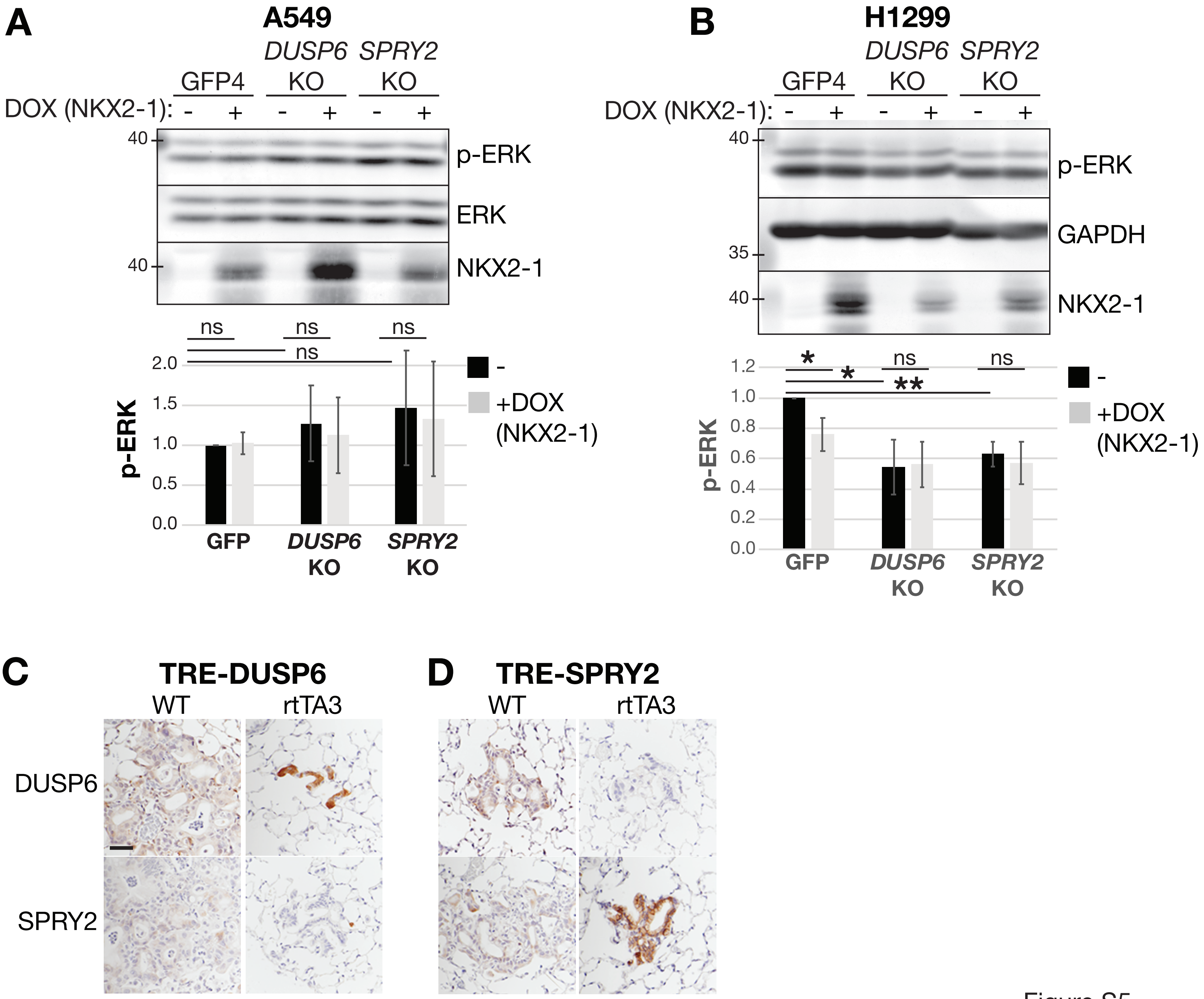
